# EMtrace: offline nanopore metagenomics on a laptop for rapid viral diagnostics, genome recovery and 16S profiling

**DOI:** 10.64898/2026.11.09.765303

**Authors:** Ilya Weinstein, Teemu Smura, Onnia Leppanen, Martyna Krupińska, Hanna Vauhkonen, Tarja Sironen, Maciej Grzybek, Olli Vapalahti, Ravi Kant

## Abstract

Nanopore sequencers produce reads within minutes, but metagenomic analysis remains challenging where internet access is limited, data are sensitive or computing resources are modest. EMtrace is an offline pipeline that runs on a laptop during sequencing: it classifies MinKNOW reads, estimates taxon abundance by expectation–maximization (EM) and assembles reference-based consensus genomes. We benchmarked it against five virome and four 16S rRNA tools on simulated, mock-community and clinical data. On field isolates and 16 SARS-CoV-2 clinical samples, EMtrace had the highest species-level F1 score (1.00 and 0.97) and one false-positive species, against 3–7 for the comparators. It recovered divergent viruses as completely as ViMOP, and low-titre field libraries more completely, at ≥99.7% identity. In replayed runs, it identified the target virus in the first 4,000-read batch (5–13 min). With a shared 16S reference, EMtrace reported two to three times fewer false-positive species than Emu.

## Main

Metagenomic sequencing can identify a pathogen without prior knowledge of what is present, and Oxford Nanopore Technologies (ONT) instruments make it portable: a MinION runs from a laptop, and reads are available while the flow cell is still sequencing^1–4^. Portable sequencing has been used in outbreak response for Ebola, Lassa and Zika viruses^3,5,6^ and in field surveys far from established laboratory settings^7^. The first real-time metagenomic analysis of clinical samples identified chikungunya and Ebola virus within minutes of sequencing^2^. A decade later, however, most published analysis pipelines still work only on a finished run.

Tools for viral metagenomics meet some of four practical requirements, but rarely all of them. The first is **offline operation**: cloud platforms such as CZ ID^8^ and the public INSaFLU-TELEVIR server^9^ need a network connection, and uploading patient data can require institutional approval. The second is **accuracy**: high precision at high recall. k-mer classifiers are fast but less precise than alignment-based methods. On a viral mock community, Kraken 2-based workflows^10^, including ONT’s EPI2ME wf-metagenomics, reported hundreds to thousands of false-positive species unless thresholds tuned to a negative control were added^11^. Viral pipelines gain precision by requiring multi-classifier agreement (TELEVIR^9^), aligning reads or contigs to large databases (CZ ID^8^, MEGAN-LR^12^) or explicitely modelling read ambiguity (metaMix^13^), often come at the cost of cloud dependence or long run times. The third is **timeliness**. ViMOP^6^, a recent offline pipeline for outbreak response, produces high-quality consensus genomes but, like tools above, starts only after sequencing ends. Real-time tools such as INSaFLU-TELEVIR’s findONTime^9^, depend on a server deployment. The fourth is **accessibility**. Graphical interfaces are offered mainly by the cloud platforms. Offline tools are almost all command-line programs; the exceptions are ONT’s EPI2ME desktop application, which runs its k-mer workflow, and MEGAN’s interactive viewer, which needs alignments prepared on the command line. A command-line-only tool is a barrier to adoption in research and clinical laboratories without dedicated bioinformatics staff.

The same four requirements apply to bacterial community profiling from full-length 16S rRNA reads, a common ONT application in clinical microbiology and field surveys^7^. Here Emu^14^ is the method against which new tools are measured^15,16^. Its expectation–maximization (EM) over full- length alignments, against a compact, non-redundant reference, gives accurate species-level abundances with far fewer false positives than k-mer classification^14,15^. It meets the accuracy requirement but not the other three: it is a command-line tool, it runs after sequencing, and it covers 16S only. A laboratory that sequences both viruses and bacterial communities therefore runs two toolchains.

Unlike the existing solutions, EMtrace is designed to address all four requirements in one tool. It answers three analysis questions: which organisms are present, in what proportions, and how much of each viral genome can be recovered. It runs offline on a laptop, without network connection or data upload. For accuracy, it extends the modular annotation strategies of our short-read virome pipeline LazypipeX^17,18^ to long reads, adding an EM abundance model that uses sub-best alignments and an automatic reference selection step for consensus assembly. For timeliness, a watch mode processes MinKNOW batch files as they are written (Fig. 1). For accessibility, one engine serves a command-line interface, the real-time monitor and a desktop graphical interface (GUI). GUI runs on the laptop that drives the sequencer, so laboratories without bioinformatics support can run the full workflow, from raw reads to abundance tables and consensus genomes.

**Figure 1.**
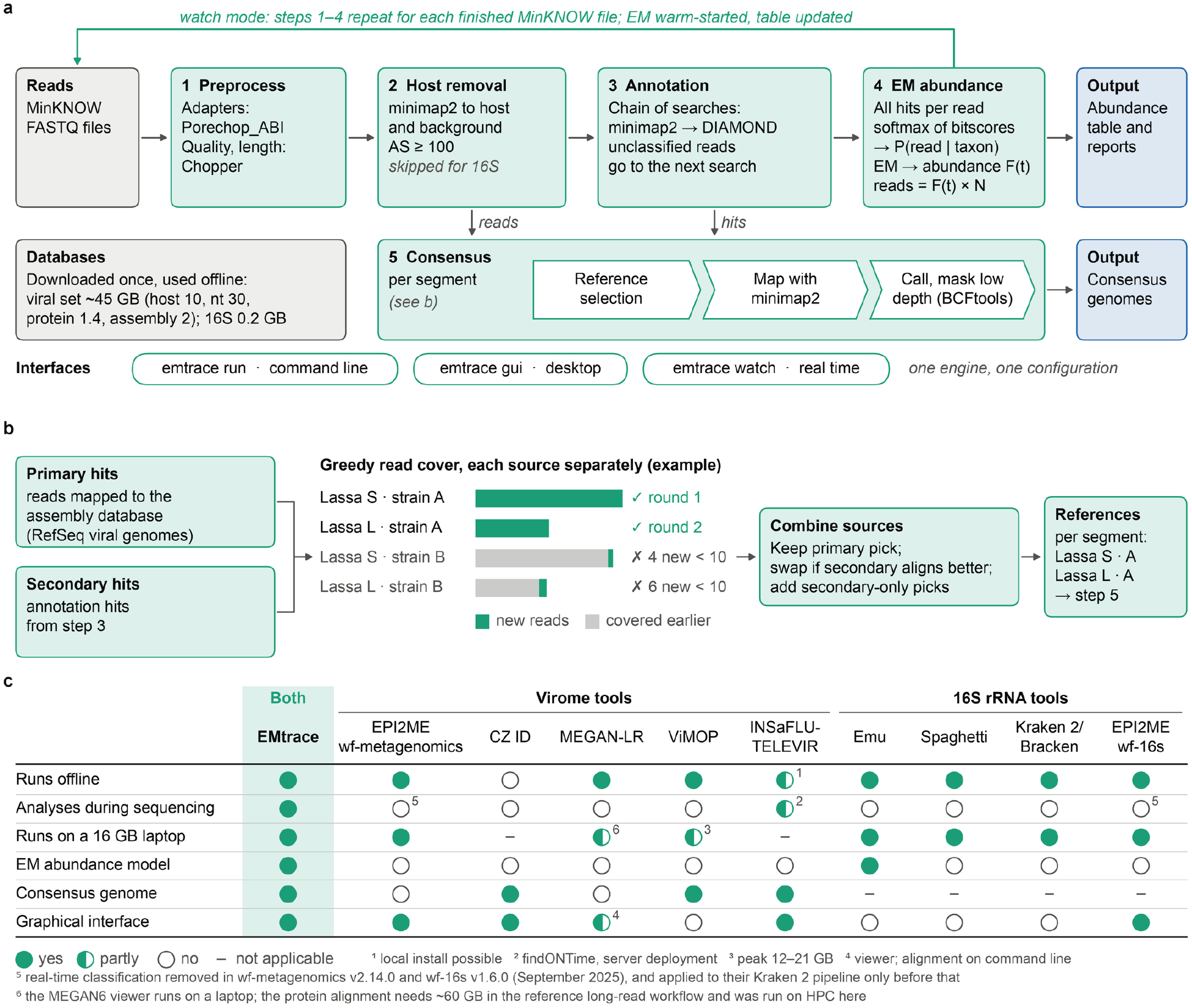
| EMtrace overview. (a) Pipeline stages and the three interfaces (CLI, GUI, watch). (b) Automatic reference selection: primary (assembly database) and secondary (annotation) hit sources, greedy read cover, and per-segment picks. (c) Operational comparison with the benchmarked tools: offline operation, processing during sequencing, laptop-class memory, abundance model, consensus assembly and graphical interface.

Benchmarking a pipeline that performs several tasks risks mixing questions that need different data. We therefore split the evaluation into separate frameworks, each with its own datasets, ground-truth and comparators (Supplementary Table 1): detection accuracy was benchmarked against EPI2ME, CZ ID and MEGAN-LR^12^; consensus assembly against ViMOP and INSaFLU- TELEVIR; 16S rRNA profiling against Emu, Spaghetti^7^, Kraken 2/Bracken^19^ and ONT’s EPI2ME wf-16s workflow^15,20^, with the reference database either chosen by each tool or held fixed. Real- time performance was evaluated by replaying sequencing runs batch by batch. Every tool was scored by the same open code, ngs-benchmark, which reads each pipeline’s native output. The scoring is therefore transparent, and every number reported here can be reproduced.

## Results

### EMtrace design

EMtrace runs five stages on each input (Fig. 1a). Reads are trimmed of adapter sequences with Porechop_ABI^21^ and filtered by quality and length with Chopper^22^. Host and background reads are removed by minimap2^23^ alignment to one or more genomes, except for 16S data. On simulated ONT data, alignment threshold of 100 removed 95.7% of human reads, no viral reads and 0.002% of bacterial reads (Supplementary Note 1). The remaining reads are classified by an *annotation strategy*: a user-defined series of minimap2, BLASTN or DIAMOND^24^ searches. In a chain, reads left unclassified by one search are passed on to a more sensitive search.

Protein searches run on open reading frames predicted by orfipy^25^. The pipeline ships curated RefSeq-, GenBank- and UniRef-derived databases that are downloaded once and used offline.

### Abundance estimation

Ambiguous reads are the main source of false positives in long-read metagenomics: a read aligning to several related species is typically assigned to the top hit, or to their lowest common ancestor. EMtrace instead keeps all alignments of each read, converts the best bitscore per read and taxon into an alignment probability with a temperature-scaled softmax^26,27^, and estimates taxon abundances *F*(*t*) by EM (Methods), shifting the weight of ambiguous reads onto taxa supported by the rest of the library. This design is similar to Emu^14^, but EMtrace accepts nucleotide or protein hits from any supported aligner, for shotgun and 16S data alike, whereas Emu uses minimap2 alignments of 16S amplicons only. The per-taxon expected read count *F*(*t*) × *N* , where *N* is the number of aligned reads, is used as the detection count throughout this study.

### Consensus assembly

When the user gives no reference, EMtrace selects one automatically (Fig. 1b). It combines two sources of hits: reads searched against a viral reference-genome database (primary) and the annotation hits (secondary). It then picks the reference genome by a greedy read cover. The reference genome that best explains the not-yet-covered reads of a species is taken, as long as it recruits at least 10 reads. Each segment of a segmented genome thus gets its own reference, while closely related strains of one segment compete for the same reads and only the best is kept. All host-filtered reads are then mapped to the chosen references with minimap2 at a sensitive seed setting (*k* = 11, *w* = 5), variants are called with BCFtools’ ONT model^28^, positions below a depth threshold are masked with N, and only the dominant allele enters the consensus (Methods).

### Real-time operation

In watch mode, EMtrace monitors a MinKNOW output directory, waits until each new FASTQ file is complete, runs it through preprocessing, host filtering and annotation, and updates the cumulative EM estimate. The estimate is warm-started from values estimated for the previous batch, to speed up convergence. The abundance table and report are updated after every file.

### Detection accuracy on clinical and field samples

We first evaluated whether EMtrace correctly identifies the viruses present in a sample. We compared it with EPI2ME wf-metagenomics (Kraken 2 + Bracken), the ONT-supported workflow; CZ ID, a widely used cloud platform; and MEGAN-LR, a long-read lowest-common- ancestor classifier. All tools were scored at species rank with the same truth and detection thresholds: ≥10 reads and ≥0.01% of the sample’s classified non-host reads (Methods).

On four single-virus field libraries from the ViMOP benchmark^6^ (Zika, West Nile, SARS-CoV-2 and dengue viruses), EMtrace identified every virus with no false positive (precision, recall and F1 all 1.00; Fig. 2a). EPI2ME and CZ ID also found every virus but reported extra species in the Zika and dengue libraries (F1 0.73 each). MEGAN-LR made no false call but missed SARS- CoV-2, assigning the few viral reads above species rank (F1 0.86).

**Figure 2.**
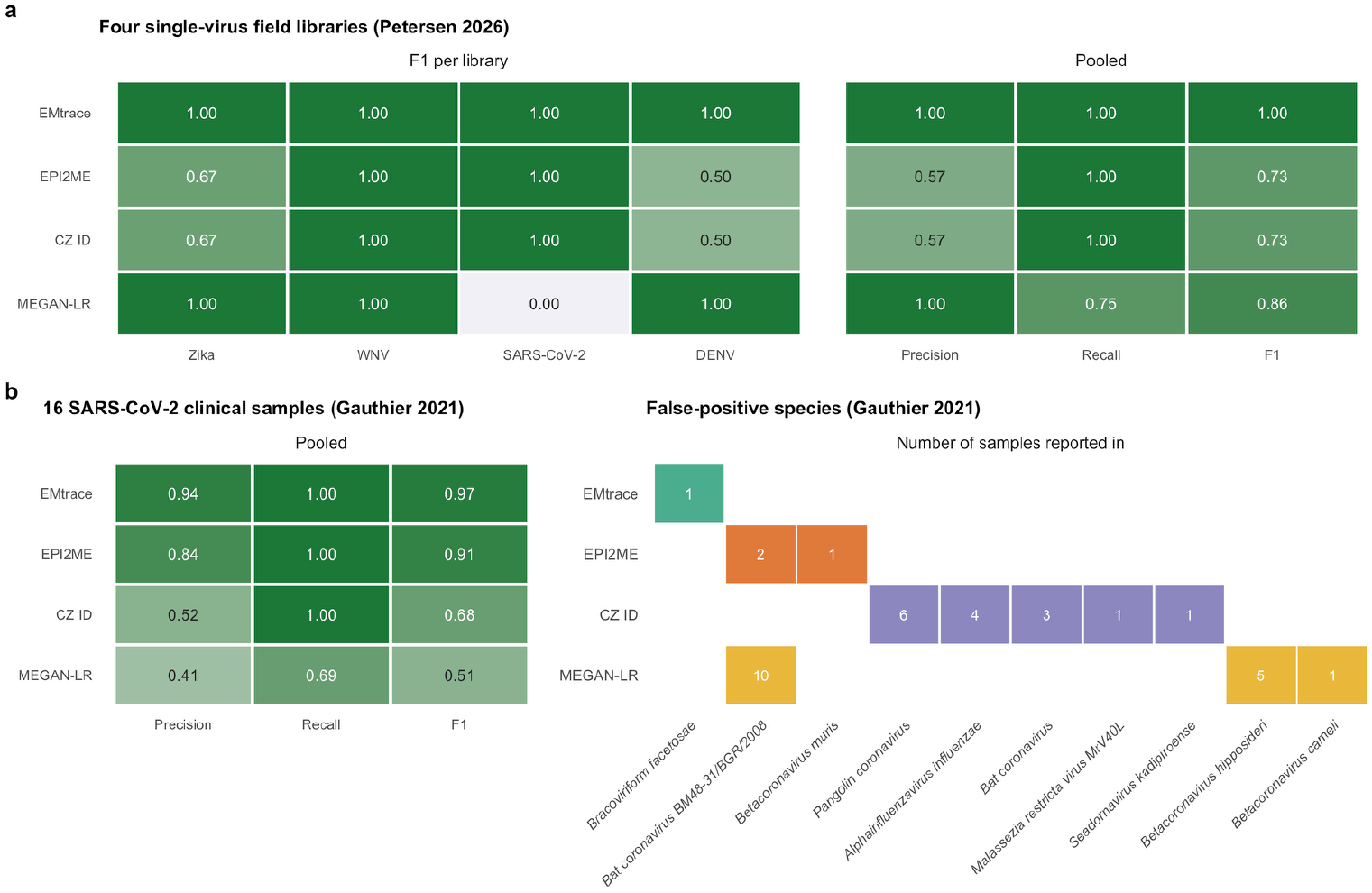
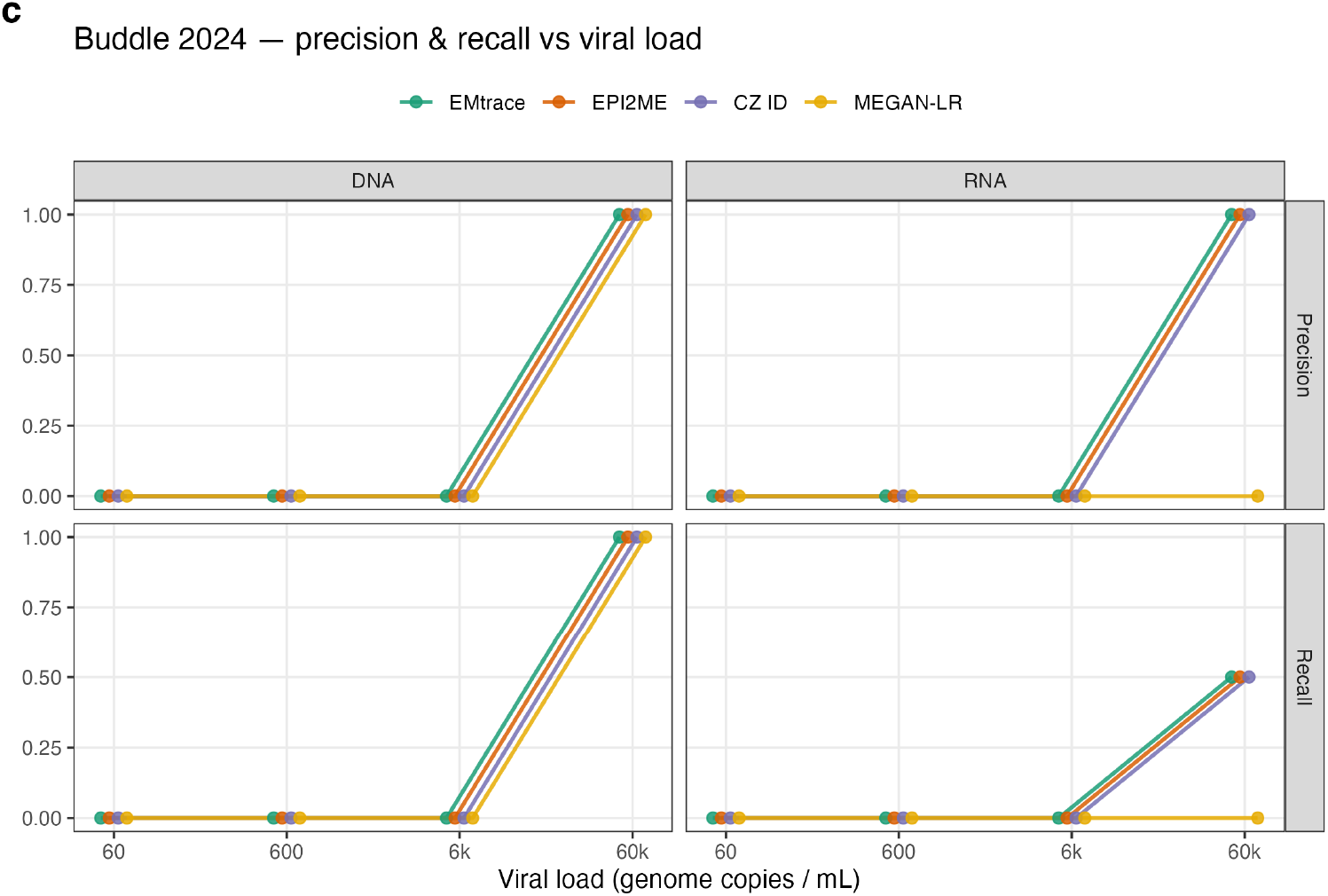
| Detection accuracy. Species rank, true and false positives pooled over the samples of each dataset. (a) Species F1 per library on the four field libraries, and pooled precision, recall and F1. (b) Pooled precision, recall and F1 per tool on 16 SARS-CoV-2 clinical samples, as in the pooled block of a (left), and the false-positive species each tool reported; the number in each tile is the number of samples in which the tool reported that species (right). (c) Species-level precision (top) and recall (bottom) against viral load for the DNA and RNA libraries of the mock community (one library per load).

Sixteen of the 20 mNGS-positive SARS-CoV-2 samples of Gauthier et al.^28^ (Methods) span RT- qPCR Ct 14.7 to 30.6. All pipelines, except MEGAN-LR, detected SARS-CoV-2 in all 16 and differed only in precision (Fig. 2b). Counting false positives as species–sample calls, EMtrace made 1 (F1 0.97), EPI2ME 3 (2 species; F1 0.91), CZ ID 15 (5 species; F1 0.68) and MEGAN-LR 16 (3 species; F1 0.51). Most of the comparators’ false positives were related coronaviruses and did not overlap with EMtrace’s. MEGAN-LR missed SARS-CoV-2 in the five samples above Ct 25.

On a six-virus mock community in human background, diluted from 60,000 to 60 genome copies per mL ^11^, EMtrace, EPI2ME and CZ ID crossed the 10-read threshold only at the highest concentration (Fig. 2c), where all four tools detected both DNA viruses, and all but MEGAN-LR detected Zika virus and respiratory syncytial virus. Below the threshold, EMtrace still found spiked viruses at the three highest concentrations (1–9 reads; Supplementary Table 2), so detection in this mock is limited by the number of viral reads, not by the classifier. We analysed one replicate library per concentration and nucleic-acid type (n = 1 per load).

### Genome recovery across divergence

Surveillance and outbreak studies often require sequencing the genome of the identified pathogen. We compared EMtrace with ViMOP^6^, which assembles by reference refinement and Medaka^29^ polishing, and INSaFLU-TELEVIR^9^, run on its public server as delivered. EMtrace maps the reads to an automatically selected reference, calls a consensus and polishes it with Medaka (Methods). Following the ViMOP benchmark design^6^, we generated our own variants at 0–30% nucleotide divergence from four target sequences: SARS-CoV-2, dengue virus 2, and Lassa virus L and S segments, with substitutions favouring sites that vary among related genomes. We then simulated ONT reads at about 30× mean depth and analysed each read set alone (“pure”) or mixed with 500,000 simulated human and bacterial background reads (“mixed”). All three tools were run on the same reads. Each consensus was aligned to the divergent genome it was simulated from. Coverage is the fraction of that genome reported as non-N bases, identity is measured over those bases, and indels are counted as alignment gaps (Methods).

Because variants at high divergence model novel viruses, we also ran EMtrace with an annotation chain that adds a protein search, which tolerates greater genetic distance than nucleotide search^8,18^. In this configuration (“chain2”), reads left unclassified by the nucleotide index are searched with DIAMOND^24^ against UniRef100 viral proteins. Chain2 had the highest recovery of the three tools at every divergence level, with a mean genome coverage of 74.9% over the 28 variants versus 64.2% for ViMOP and 52.8% for TELEVIR (Fig. 3a and Supplementary Table 3). It chose the correct reference in all 27 variants that gave a consensus and made no off-target selection. The only failure, the Lassa L segment at 30% divergence, was not usefully recovered by any tool (ViMOP 4.7% coverage).

**Figure 3.**
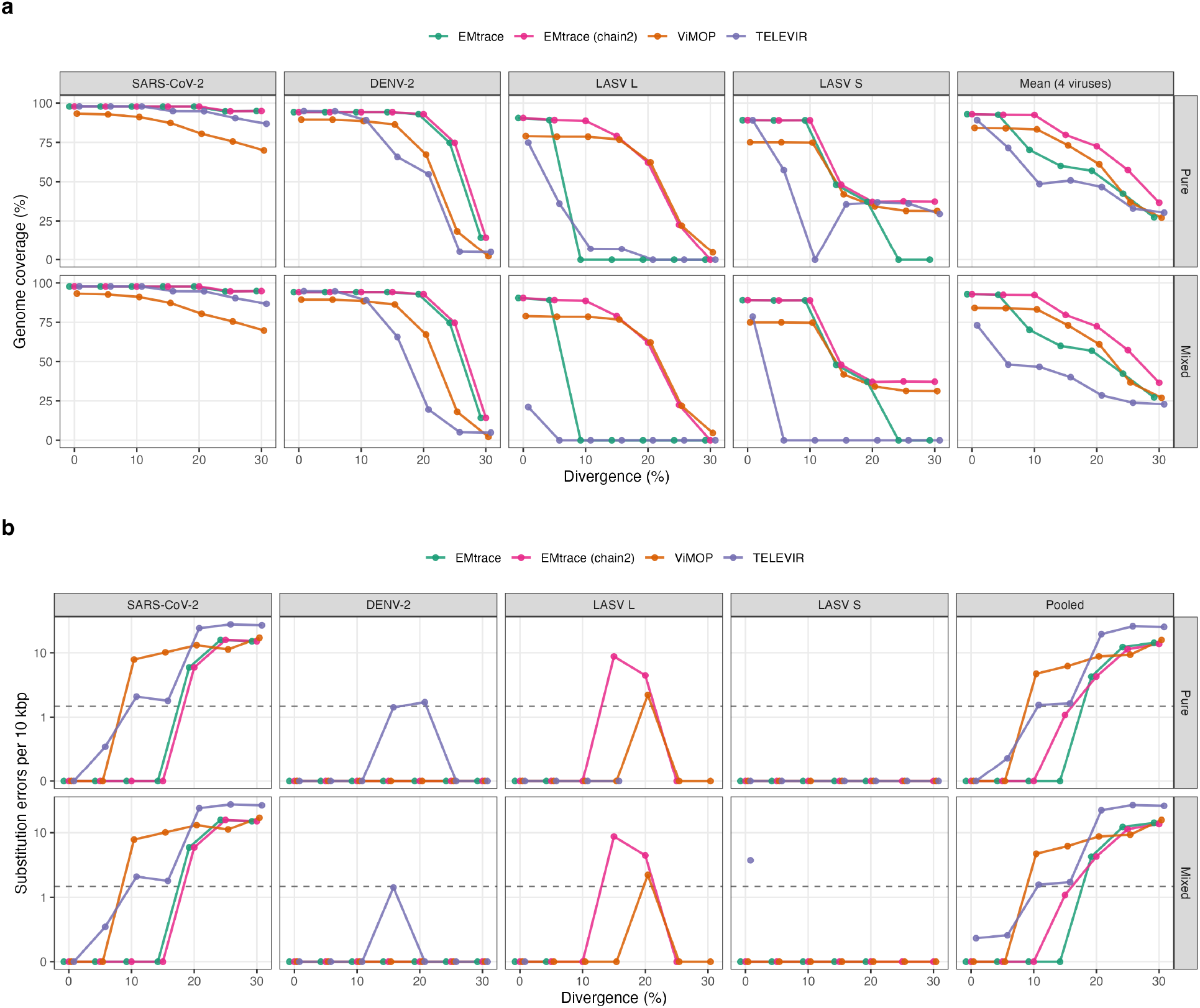
| Genome recovery across divergence. (a) Genome coverage against divergence from the source genome, per virus and averaged, for pure (top) and background-embedded (bottom) reads. (b) Substitution errors per 10 kbp of aligned consensus (log scale). The dashed line marks the error expected from majority calling at 10× depth and 10% per-read error. DENV-2, dengue virus 2; LASV L and S, Lassa virus L and S segments. Background reads did not change EMtrace’s results in any of the 28 pure–mixed pairs, because the background adds no reads that map to the viral reference. ViMOP was similarly unaffected. For TELEVIR, background reduced mean coverage from 52.8% to 40.5% (Supplementary Table 3).

With the default nucleotide annotation, EMtrace achieved the highest genome coverage at 0– 5% divergence but trailed ViMOP at 10–20%, for a mean of 63.2% over the 28 variants (Supplementary Table 3). The deficit was entirely due to Lassa virus: SARS-CoV-2 and dengue virus were recovered better than by ViMOP at every divergence level, and identically to chain2. For Lassa L at 10–30% and Lassa S at 25–30% divergence, the nucleotide index classified too few reads for automatic selection to pick a reference, so there was no consensus. Chain2 recovered six of these seven variants, raising mean Lassa coverage from 38.0% to 61.3% (ViMOP 54.7%).

At low divergence, the tools recovered similar fractions of each genome, with EMtrace slightly ahead. At higher divergence, performance differed by virus: EMtrace recovered more of SARS- CoV-2 and dengue virus, whereas on Lassa L chain2 and ViMOP performed similarly (Fig. 3a).

The consensus bases that every tool reported were accurate; EMtrace had the lowest error rate. Pooled over all pure simulated samples, EMtrace miscalled 3.6 bases per 10 kbp with both default and chain2 settings, versus ViMOP 5.3 and TELEVIR 8.6 (Fig. 3b). Up to 15% divergence, EMtrace’s default consensus contained no miscalled bases, whereas ViMOP had 4.7 and 6.2 per 10 kbp at 10% and 15%, respectively. The remaining errors lay in SARS-CoV-2 at ≥20% divergence, where unresolved variant sites keep the reference base; here EMtrace and ViMOP converged (4–16 per 10 kbp), whereas TELEVIR made more errors (19–25). EMtrace also made the fewest indels, the predominant error type in nanopore consensus sequences: 1.9 per 10 kbp versus 3.7 for ViMOP and 3.9 for TELEVIR (Supplementary Table 3).

### Genome recovery from real libraries

On the four field libraries (Supplementary Fig. 1a; scoring in Methods), EMtrace and ViMOP performed similarly on the two high-load libraries, Zika and dengue virus (≥99.3% coverage, ≥99.2% identity), whereas TELEVIR mapped the dengue virus 2 reads to a dengue virus 1 reference (83.8% coverage). On the low-titre West Nile virus library, EMtrace achieved the highest genome coverage (42.2%, against 32.2% for TELEVIR and 25.6% for ViMOP). The SARS-CoV-2 library (about 365 viral reads) was depth-limited for every tool (at most 29.3% coverage). Overall, EMtrace had the highest mean coverage (65.5%, against 61.2% for TELEVIR and 56.1% for ViMOP) at the highest mean identity (99.7%, equal to ViMOP).

EMtrace and ViMOP each made a single indel, both on West Nile virus (0.3 and 0.4 per 10 kb), whereas TELEVIR made 5.3 indels per 10 kb.

In the 16 Gauthier clinical samples, recovery depended sharply on viral load, with a cliff at Ct ≈ 25 (Supplementary Fig. 1b). All three tools produced near-complete genomes (≥99% coverage) from the 11 samples at Ct ≤ 25.4, and all recovered less than 20% from the five samples at Ct ≥ 25.5, where the number of viral reads fell from 4,539 to 39. Mean coverage was 70.7% for TELEVIR, 70.0% for EMtrace and 68.8% for ViMOP. Identity to Wuhan-Hu-1 was ≥99.7% on all near-complete genomes. Because the samples belong to several lineages and no sample- specific reference genome is available (Methods), this identity includes lineage differences.

### Full-length 16S rRNA profiling with the reference held fixed

Emu showed that EM over full-length ONT 16S alignments gives accurate species profiles^14^, and it is the natural reference for EMtrace’s abundance model. Comparing the two raised a methodological problem common to many classifier benchmarks. Each tool ships with its own reference database, which can matter more than the algorithm: on a 21-species synthetic gut community^14^, EMtrace’s genus L1 error was 0.351 on its own 16S database and 0.122 on Emu’s. We therefore report two comparisons: each tool on its own database, which is what a user gets by default (Supplementary Table 4), and every tool on Emu’s 49,243-sequence reference, which isolates the method (Table 1).

**Table 1.**
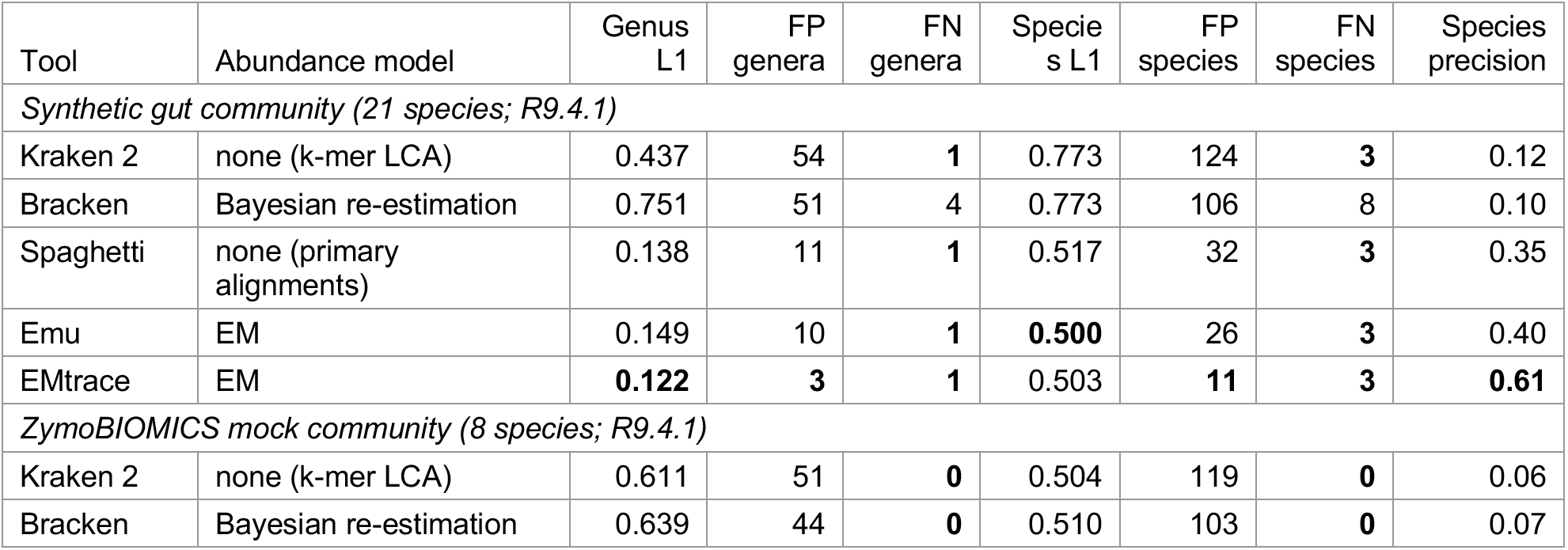
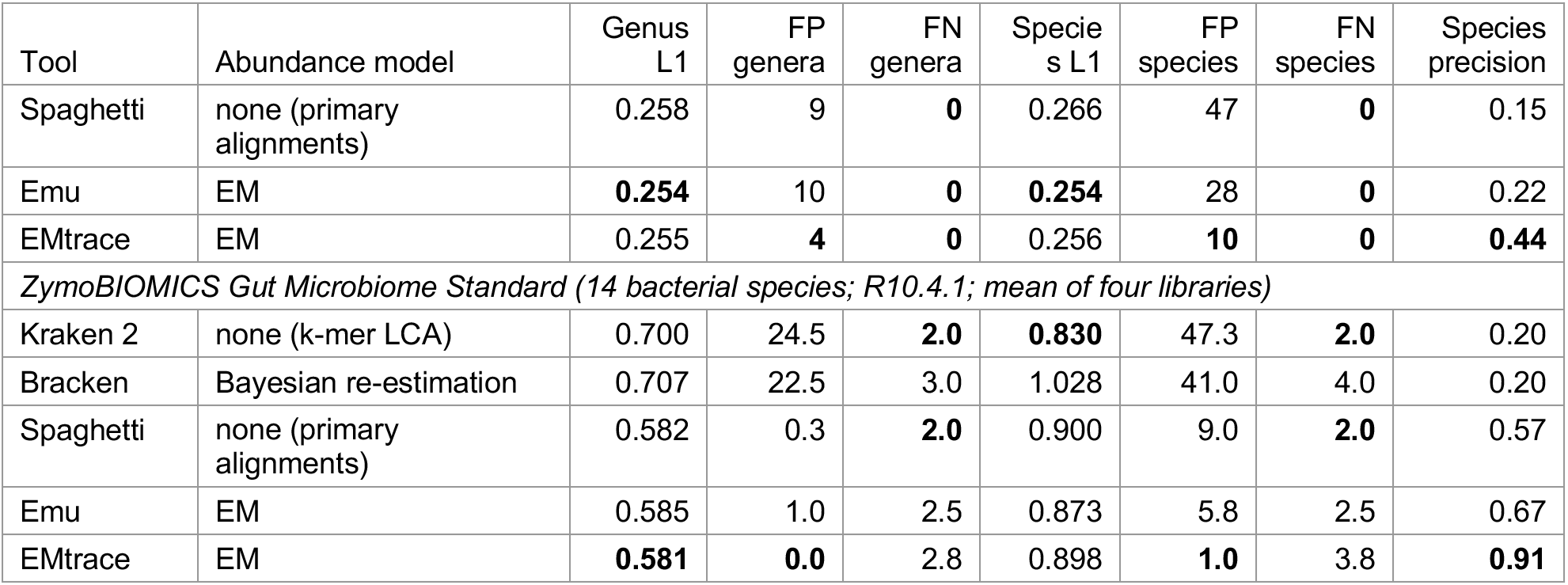
| 16S profiling with all tools on one reference. All tools on Emu’s 49,243-sequence 16S database, synonyms collapsed, with a 10-read threshold and a 0.01% abundance floor. Synthetic gut community and ZymoBIOMICS, one library each; Gut Microbiome Standard, mean of four libraries (s.d. of L1 ≤0.005). L1, sum of absolute differences between predicted and true relative abundances (0–2; lower is better). EMtrace and Emu report EM estimates, the other tools read counts. FP, reported taxa absent from the truth; FN, truth taxa not reported. Bold, best per column within each community.

| Tool | Abundance model | Genus L1 | FP genera | FN genera | Species L1 | FP species | FN species | Species precision |
|---|---|---|---|---|---|---|---|---|
| *Synthetic gut community (21 species; R9.4.1)* |  |  |  |  |  |  |  |  |
| Kraken 2 | none (k-mer LCA) | 0.437 | 54 | **1** | 0.773 | 124 | **3** | 0.12 |
| Bracken | Bayesian re-estimation | 0.751 | 51 | 4 | 0.773 | 106 | 8 | 0.10 |
| Spaghetti | none (primary alignments) | 0.138 | 11 | **1** | 0.517 | 32 | **3** | 0.35 |
| Emu | EM | 0.149 | 10 | **1** | **0.500** | 26 | **3** | 0.40 |
| EMtrace | EM | **0.122** | **3** | **1** | 0.503 | **11** | **3** | **0.61** |
| *ZymoBIOMICS mock community (8 species; R9.4.1)* |  |  |  |  |  |  |  |  |
| Kraken 2 | none (k-mer LCA) | 0.611 | 51 | **0** | 0.504 | 119 | **0** | 0.06 |
| Bracken | Bayesian re-estimation | 0.639 | 44 | **0** | 0.510 | 103 | **0** | 0.07 |
| Tool | Abundance model | Genus L1 | FP genera | FN genera | Species L1 | FP species | FN species | Species precision |
| Spaghetti | none (primary alignments) | 0.258 | 9 | **0** | 0.266 | 47 | **0** | 0.15 |
| Emu | EM | **0.254** | 10 | **0** | **0.254** | 28 | **0** | 0.22 |
| EMtrace | EM | 0.255 | **4** | **0** | 0.256 | **10** | **0** | **0.44** |
| *ZymoBIOMICS Gut Microbiome Standard (14 bacterial species; R10.4.1; mean of four libraries)* |  |  |  |  |  |  |  |  |
| Kraken 2 | none (k-mer LCA) | 0.700 | 24.5 | **2.0** | **0.830** | 47.3 | **2.0** | 0.20 |
| Bracken | Bayesian re-estimation | 0.707 | 22.5 | 3.0 | 1.028 | 41.0 | 4.0 | 0.20 |
| Spaghetti | none (primary alignments) | 0.582 | 0.3 | **2.0** | 0.900 | 9.0 | **2.0** | 0.57 |
| Emu | EM | 0.585 | 1.0 | 2.5 | 0.873 | 5.8 | 2.5 | 0.67 |
| EMtrace | EM | **0.581** | **0.0** | 2.8 | 0.898 | **1.0** | 3.8 | **0.91** |

We used three mock communities (Methods): the ZymoBIOMICS Microbial Community Standard (8 species; R9.4.1), the synthetic gut community^14^ (21 species; R9.4.1), and the same manufacturer’s Gut Microbiome Standard (15 species; R10.4.1)^30^.

With the reference held fixed, false positives accounted for most performance differences among the five tools (Table 1). False negatives were few and the same for all tools except Bracken: on the synthetic gut community, one missed genus and three missed species (Bracken 4 and 8), and none on the ZymoBIOMICS mock (Table 1). The missed taxa were either absent from the shared reference (*Romboutsia hominis*) or below the 10-read/0.01% threshold (e.g. *Gemmiger formicilis*).

On the synthetic gut community, EMtrace had the lowest genus L1 error (0.122; Emu 0.149, Spaghetti 0.138) and 3 false-positive genera (Emu 10, Spaghetti 11). At species rank, EMtrace and Emu had similar L1 (0.503 and 0.500), but EMtrace reported 11 false-positive species against Emu’s 26, giving 1.5-fold higher precision (0.61 versus 0.40). Both assigned about 20% of the reads outside the ground truth, mostly those of the absent *R. hominis*.

The ZymoBIOMICS mock showed a similar pattern: the two tools tied on L1 at both ranks (EMtrace 0.255 and 0.256, Emu 0.254), and EMtrace reported 10 false-positive species against 28 and 4 false-positive genera against 10 for Emu (Table 1).

The controlled comparison separates two properties that are often reported as one: accurate taxon proportions and the correct set of taxa. Spaghetti, which counts primary alignments without abundance model, matched the EM tools on genus L1 and missed no more genera, but reported 11 false-positive genera against 3 for EMtrace. The EM was therefore not needed for accurate proportions; its contribution was fewer false-positive calls. The k-mer tools performed worse on both counts: on the same reference, Kraken 2 had 3.6 times EMtrace’s genus L1 and 11 times as many false-positive species. Bracken re-estimation shortened the false-positive list but also removed true-positive genera, raising genus L1 to 0.751. As Emu also uses EM, EMtrace’s higher precision is most likely explained by the information supplied to the model: it considers more secondary alignments per read and weighs near-best hits by a temperature- scaled softmax.

On their own databases (same thresholds; Supplementary Table 4), EMtrace (de-duplicated rrnDB^31^) led Emu at species rank on the synthetic gut community (L1 0.446 against 0.500; 12 against 26 false-positive species) but trailed at genus rank (0.351 against 0.149): rrnDB has no named *Romboutsia*, so the *R. hominis* reads were assigned to *Intestinibacter*. The error disappeared on Emu’s reference, which contains the genus.

To test current chemistry and one more community, we used ONT’s public R10.4.1 runs of the ZymoBIOMICS Gut Microbiome Standard^30^: four replicate libraries of 14 bacterial species and one archaeon (Methods). The replicates agreed closely (L1 SD≤0.005). Every alignment-based tool had a genus L1 of about 0.58 with almost no reads outside the ground-truth genera: all over-called *Bacteroides* and *Veillonella* and under- called *Limosilactobacillus*, *Roseburia* and *Bifidobacterium*, an error most likely from the dataset, not the classifiers. On Emu’s reference the tools again differed in the taxon list: EMtrace reported no false-positive genus and one false- positive species (species precision 0.91), against one genus and 5.8 species for Emu (0.67) and 0.3 genera and 9.0 species for Spaghetti (0.57), although Emu had a lower species L1 (0.873 against 0.898; Table 1). Every tool assigned most of the *Veillonella rogosae* reads (15.9% of the community) to *V. parvula*, whose 16S sequence differs at about 1% of positions. Tools that spread these reads over several *Veillonella* species recovered part of the *V. rogosae* abundance: Kraken 2 had the lowest species L1 on this reference (0.830) but 47 false-positive species.

On its own database (Supplementary Table 4), EMtrace assigned these reads to *V. rogosae* and had the lowest species L1 of all tools and databases (0.581), with no false-positive species and no reads outside the ground-truth. ONT’s wf-16s performed similarly at genus rank (L1 0.584) but not at species rank (L1 1.359). Every tool missed the taxa below the 10-read threshold by design (*Enterococcus faecalis*, *Clostridium perfringens*) and the archaeon *Methanobrevibacter smithii*; *Salmonella enterica* fell below the threshold in some libraries for both EMtrace and Emu.

### Real-time detection during a sequencing run

Among the benchmarked tools, only EMtrace reports during the run on the laptop that drives the sequencer. We replayed four viral libraries (two in-house runs and two public tick pools; Methods), one 16S MinKNOW run file by file, and four R10.4.1 replicate libraries of a 16S gut mock through emtrace watch on a laptop, scoring every batch against the truth or, for the tick pools, against the published findings (Methods). Two clocks are reported: *sequencing time*, when an answer became possible, and *compute time*, how long EMtrace took to produce it; the public tick-pool reads carry no start times, so only compute time is reported for them.

Kevo orbivirus (KEVOOV; isolate fi663), an untargeted cell-culture library^32^, is absent from the annotation databases (nearest relatives ∼71% nucleotide identity) and was scored at genus rank (*Orbivirus*). Tick-borne encephalitis virus (TBEV; four pooled tick samples, a virus-specific amplicon library; Methods) is in the databases and was scored at species rank. The two tick pools, 58-37 and 35-33, are public untargeted libraries of field-collected *Rhipicephalus bursa*^33^ with no ground truth, scored instead against the published findings: Crimean–Congo haemorrhagic fever virus (CCHFV, *Orthonairovirus*) and Jingmen tick virus (JMTV, *Jingmenvirus*) by in-house PCR, and the viruses reported from nanopore sequencing alone (Methods). KEVOOV and TBEV topped the first 4,000-read batch and every later one (Table 2), which closed 13.1 and 4.8 min into sequencing and was reported 3.8 and 1.4 min later. The target’s EM probability never fell below 0.9995 for TBEV and 0.97 for *Orbivirus* in KEVOOV (summed over its database relatives), and other taxa stayed minor (≤810 reads and EM probability ≤0.006 for KEVOOV; for TBEV, only Louping ill virus reached 10 reads). In both pools, every PCR-positive virus reached 10 reads in the first batch and stayed there; the PCR- negative JMTV of pool 35-33 never did (2 reads). Final CCHFV counts (1,974 and 856) and pool 58-37’s JMTV count (3,217) were 7–13% above those published (1,752, 783 and 3,020).

**Table 2.** | Replayed sequencing runs. First correct call: the first batch from which the top-ranked taxon stays correct. Viral libraries were annotated with DIAMOND added, except TBEV (minimap2 only; Methods). Viral reads: reads aligned to the target genome(s) in the final assembly, as a percentage of the library (pool 58-37: CCHFV / JMTV). *Tick pools: first batch from which every PCR-positive virus has ≥10 reads; no read start times, so no sequencing time or throughput. 16S mock: first batch from which all eight truth genera and species are recovered with no false positive. Throughput ratio: sequencing time / compute time. Gut mock: four R10.4.1 replicates, re-cut into 4,000-read files by read start time; ranges over replicates. No batch recovered all 15 truth taxa; four lie below the read threshold by design.

| Library | Viral reads (%) | Batches | Run length | First correct call: batch/ sequencing time / reads | Compute per batch (median) | Throughput ratio |
|---|---|---|---|---|---|---|
| KEVOOV, isolate fi663 | 99.0 | 116 | 19.0 h | 1 / 13.1 min / 4,000 | 271 s | 2.1 |
| TBEV, 4 pooled tick samples | 46.5 | 79 | 5.0 h | 1 / 4.8 min / 4,000 | 93 s | 2.4 |
| Tick pool 58-37 | 5.0 / 15.8 | 12 | — | 1* / — / 4,000 | 152 s | — |
| Tick pool 35-33 | 0.39 | 55 | — | 1* / — / 4,000 | 136 s | — |
| barcode79 (16S mock) | — | 26 | 71.8 h | 1 / 85.6 min / 4,000 | 294 s | 34.9 |
| Gut mock (16S, 4 replicates) | — | 19–22 | 1.9 h | — (L1 stable by batch 1–6, ≤39 min) | 183 s | 1.9 (1.8–2.1) |

EMtrace kept pace with sequencing on every library with read start times, 1.8–34.9 times faster than reads were produced (Table 2). A 4,000-read viral batch took a median of 93–271 s, about a minute of it loading the host and annotation indexes; DIAMOND raised the KEVOOV median from 184 to 271 s. The protein search mattered most for divergent viruses: DIAMOND raised KEVOOV’s *Orbivirus* reads from 49,334 to 128,075 and pool 58-37’s JMTV from 738 to 3,217 (Fig. 4a). Only with DIAMOND did a *Mivirus* (both pools) and a *Phlebovirus* (58-37), both reported in the original study from nanopore sequencing alone, reach 10 reads. For CCHFV, with closer relatives in RefSeq, it added only 5–14%.

**Figure 4.**
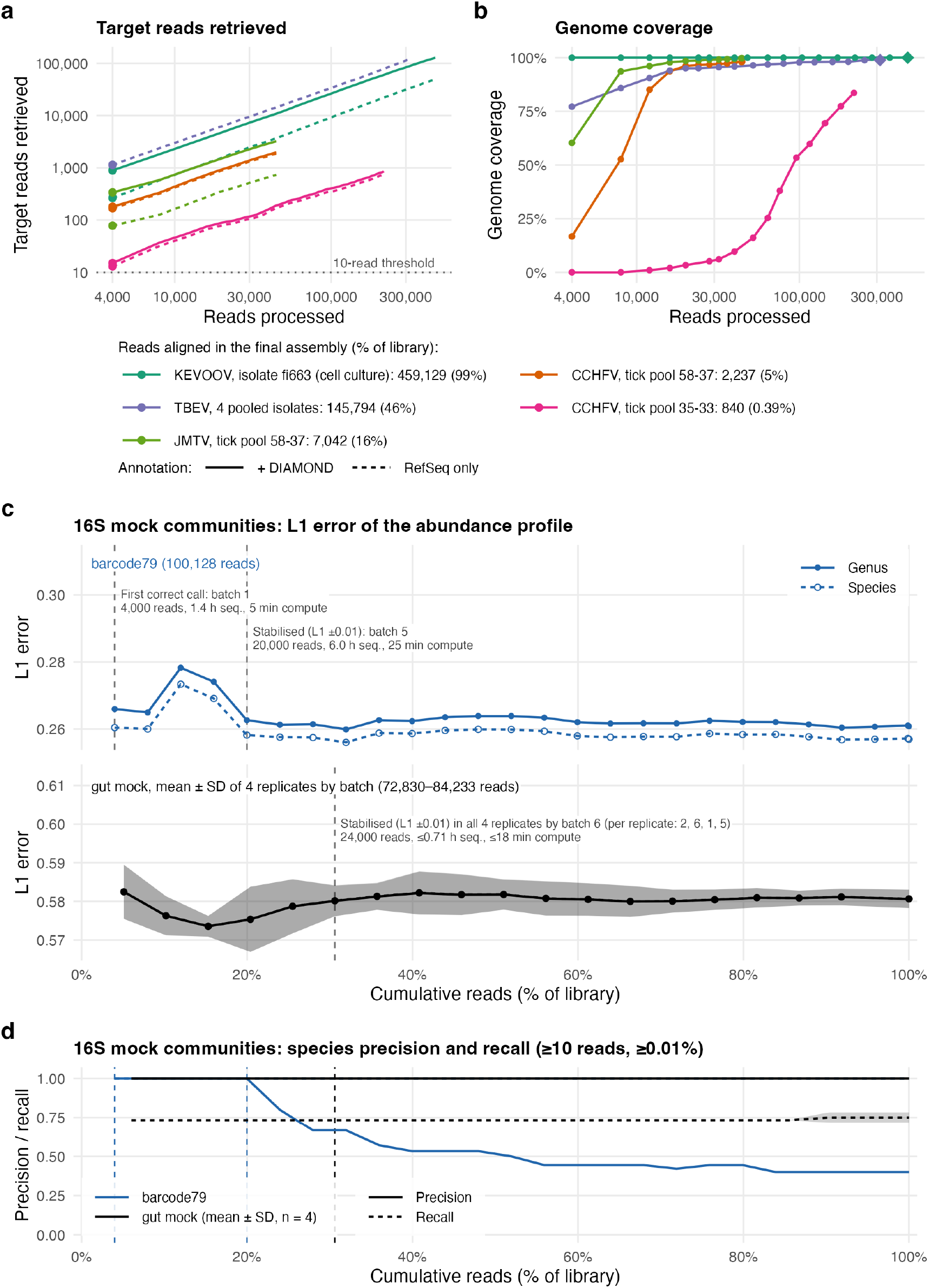
| Real-time replay. (a) Target-virus reads retrieved against reads processed (both log scale) for the four viral libraries; solid, with DIAMOND; dashed, minimap2 against RefSeq only (TBEV was annotated with RefSeq only); points mark the first 4,000-read batch. (b) Genome coverage of the consensus assembled against a reference genome of each target from the reads of batches 1…k (log- spaced k, minimum depth 10); diamonds, final assembly polished with Medaka. The legend gives the reads aligned in the final assembly and their percentage of the library. Tick pools are named by the pool IDs of the original study (BioProject PRJNA878809; pool 58-37: SRR21504617, pool 35-33: SRR21504592). Panels c and d are plotted against cumulative reads as a percentage of each library sequenced (library sizes in the legends). (c) 16S mock communities: genus (solid) and species (dashed) L1 error per batch for barcode79 (top) and the ZymoBIOMICS Gut Microbiome Standard (bottom; four R10.4.1 replicates, mean ± SD); for the gut mock the genus and species curves coincide. Dashed vertical lines mark the first correct call and stabilisation (genus and species L1 within ±0.01 of their final values; gut mock, the latest of the four replicates). (d) Species precision and recall for the same libraries; vertical lines as in c. barcode79 recall stays at 1.00, hidden under the gut-mock precision line. The gut-mock batches were re-cut from the read start times at 4,000 reads per file, because the public data hold no MinKNOW batch files.

To see when a usable genome becomes available, we assembled consensus genomes from the reads accumulated up to log-spaced batches (Methods). KEVOOV, against its own published genome, had complete coverage of all segments by the first batch, 13.1 min into the run (Fig. 4b). TBEV covered 77.2% of Neudoerfl in the first batch, 95.1% after five (20,000 reads, 18.4 min) and 99.2% at the end; its 97.6% identity reflects the samples’ divergence from Neudoerfl genome. In pool 58-37, JMTV and CCHFV passed 90% coverage after 8,000 and 16,000 reads; in pool 35-33, with 0.39% CCHFV reads, coverage passed 50% only after about 96,000 reads and reached 83.6%. Genome recovery therefore depended on viral load, whereas detection did not: CCHFV was detected in the first 4,000 reads of both pools.

For barcode79, the question is when the *proportions* converge rather than when a taxon is first detected. The first batch, 4,000 reads and 1.4 h into the run, recovered all eight genera and species with no false positive, at L1 errors of 0.266 and 0.260 against final values of 0.261 and 0.257 (Fig. 4c); the remaining 96,000 reads changed L1 by 0.005. As sequencing continued, false-positive taxa accumulated above the 10-read threshold and species precision fell from 1.00 to 0.40 (Fig. 4d). The last batch matched a single-pass analysis of the whole library, with the same taxa above the threshold, the same precision (0.44 and 0.40) and L1 within 0.0015, so warm-started batch processing costs no accuracy.

The four R10.4.1 replicates of the ZymoBIOMICS Gut Microbiome Standard were sequenced about 30 times faster than barcode79 (about 40,000 reads per hour against 1,400), and the first batch closed 8.7–9.4 min into each run. Its genus L1 (0.578–0.593) was already within 0.012 of the final value (0.578–0.583, the static result in Supplementary Table 4), and every replicate was stable within ±0.01 by batch 6, at most 39 min in (Fig. 4c). No replicate reported a false- positive genus or species at any batch, so precision stayed at 1.00; recall stayed at 0.73–0.80, the missing genera being those below the read threshold in the static analysis (Fig. 4d).

Processing took 181–185 s per batch, 1.8–2.1 times faster than reads were generated. On current chemistry the real-time margin is therefore smaller, but EMtrace still kept pace with sequencing.

EMtrace answered while the sequencer was still running, and the first answer was the final one. Virus calls came 6 min (TBEV) and 17 min (KEVOOV) into sequencing and never changed, whereas a tool that starts at the end of the run could not begin for 5.0 and 19.0 h; in the tick pools every PCR-confirmed virus was found within the first 4,000 reads, 2–9% of each library.

The 16S profiles were available 1.4 h into the 71.8 h barcode79 run and stable within 39 min of the 1.9 h gut-mock runs — all offline, on the laptop that drives the sequencer.

## Discussion

EMtrace brings classification, abundance estimation and consensus assembly into one offline tool that runs on a laptop and reports while the sequencer is still running. In our benchmarks, this did not come at the cost of accuracy. It had the highest detection F1 on both the field libraries (1.00, the only tool with neither a false positive nor a missed virus) and the clinical samples (0.97). It matched or exceeded ViMOP on genome recovery, with the most accurate consensus sequences, and detected the target virus in the first 4,000-read batch of every replayed viral library.

A key advance of the framework is that actionable pathogen identification becomes available while sequencing is ongoing and without network connectivity. Every known or PCR-confirmed virus in the replayed viral libraries was called in the first 4,000-read batch, 6–17 min into sequencing where run times are known, and at as little as 0.4% of the reads. The time to answer was therefore set by the sequencer, not by the analysis. The gut-mock 16S profiles were near their final values from the first batch and stable within 39 min, whereas a post-run tool could only have started after about 2 h. All replays ran on a 16 GB laptop, the machine that drives the sequencer, and no data left it. This changes the workflow in three ways. First, the operator can stop the run or start targeted follow-up once the call is made (unless a whole genome is needed). Second, patient data do not have to be uploaded, so institutional approval for cloud processing is not needed. Third, a MinION and a laptop are enough to run the whole analysis, which suits the outbreak and field settings.

The most consistent result across frameworks is high precision without loss of recall. On the field and clinical datasets, EMtrace matched the best comparator’s recall (1.00) and reported fewer false positives. Virus detection on clinical samples, 16S profiling on a fixed reference, and real-time calling all show the same pattern: EMtrace’s error is concentrated on few taxa, whereas the comparators spread it over many small ones. We attribute this to the EM receiving near-best alternatives rather than top hits alone. Reads shared between related species are resolved by the evidence from the rest of the sample, instead of each creating a low-count bin. For diagnostic use, a short list with one false positive is far easier to act on than a long tail of low-count taxa, each needing confirmation by an independent method such as qPCR. Tools that tied on L1 differed two- to sixfold in false positives, so we recommend reporting false-positive counts beside any abundance error.

The 16S results held across sequencing chemistries. On two mock communities sequenced with older chemistry and on R10.4.1 replicate libraries of a third, EMtrace matched or outperformed Emu on genus-level L1 using the shared reference, came within 0.03 of it on species-level L1, and reported two to six times fewer false-positive species.

Controlling the reference database changed the conclusions of the 16S comparison, and it should also be considered for the virome results. A tool compared on its own database is compared as a product, and a database gap can look like a failure of the method, as the *Romboutsia* case showed. In the assembly benchmark, the same effect favoured ViMOP, whose reference set contains the Lassa templates from which the divergent genomes were simulated. EMtrace’s RefSeq references were already 13–16% away before any mutation was added. Even so, with protein annotation and polishing EMtrace recovered as much of Lassa L at 15–25% divergence as ViMOP.

Several limitations remain. First, Medaka polishing needs a basecaller-matched model, which public archives often omit from read headers, and on a very low-depth library it slightly reduced coverage (SARS-CoV-2 field library, 21.4% to 20.4%). Second, divergent viruses need a protein annotation step, a configuration choice users must anticipate: the default nucleotide index lost Lassa virus beyond 10% divergence. Third, the real-time results come from replays of complete runs, which reproduce the file timing and content of a live run but not the load of concurrent basecalling (on GPU-equipped laptops mainly memory, on CPU-only machines also processor time); validation during live runs is under way. The replays nevertheless left a 2.1- to 2.4-fold processing margin on the two virus libraries with read start times, and about one minute of each batch is spent reloading the host and annotation indexes, which future releases may keep in memory. Fourth, several datasets are small (four field libraries, one viral mock library per load and nucleic-acid type, and three 16S mock communities), so results are reported descriptively, without statistical tests. Finally, we did not evaluate INSaFLU-TELEVIR’s live-run module findONTime^9^, which needs an INSaFLU-TELEVIR server, whereas our benchmark tests offline analysis on the sequencing laptop.

EMtrace thus brings the full analysis onto the laptop that drives the sequencer, offline and in real time, without losing accuracy. A sequencer and a laptop are enough to go from sample to actionable result, wherever sequencing happens. EMtrace and its benchmark are open, so the comparisons can be rerun as the tools change.

## Methods

### EMtrace implementation

EMtrace (v1.0.0) is written in Python and uses external bioinformatic tools. Porechop_ABI21 trims adapters, and Chopper22 filters reads by mean quality and length. Host and background reads are removed by minimap2 alignment (map-ont preset) to one or more genomes, discarding reads with alignment score ≥100. Annotation strategies are defined in YAML. Each search is a minimap2, BLASTN, DIAMOND BLASTP (on orfipy ORFs) or DIAMOND BLASTX step against a registered database, targeting either all reads or those left unannotated by the previous step (chain). Hits are converted to a common 18-field annotation table and assigned an NCBI division (or SILVA domain). Reads are then partitioned into viral, phage, bacterial and unannotated sets. EMtrace uses SILVA taxonomy for SILVA, and NCBI taxonomy for all other reference databases. Outputs are an annotation table, an abundance table, HTML reports, and per-division reads in FASTQ format.

### EM abundance estimation

For each read *q* and taxon *t*, the best bitscore *b*(*q*, *t*) over all alignments is scaled by a temperature *T* and converted to an alignment probability by a softmax over the read’s candidate taxa, *P*(*q* | *t*) = exp(*b*(*q*, *t*)/*T*) / Σ*t*′ exp(*b*(*q*, *t*′)/*T*), following ref. ^26^.

Abundances are estimated by EM. The E-step computes *P*(*t* | *q*) ∝ *P*(*q* | *t*)*F*(*t*), and the M-step sets *F*(*t*) = Σ*q P*(*t* | *q*)*P*(*q*) with a uniform read prior. Iteration stops when the log-likelihood improves by less than 0.01 (at most 1,000 iterations). All computations are done in log space. The expected read count *F*(*t*) × *N*, where *N* is the number of reads entering the EM, is reported as readn_em and used for detection thresholds. Top-hit read counts (readn) are reported beside it. Reads whose top hits tie across taxa are credited to the tied taxon with the highest *F*(*t*). By default, all alignments up to the aligner’s secondary limit (minimap2 -N) enter the EM; --tophits restricts it to top-scoring hits. In watch mode, *F* is initialized from the previous batch’s estimate.

### Automatic reference selection

Candidate references come from two sources: the viral reads mapped to a viral genome database (RefSeq viral genomes), and the viral annotation hits. In each source, hits are grouped by species, and species with at least 10 reads (min_reads) are kept. Accessions are then chosen by greedy cover: each round takes the accession with the highest summed alignment quality (matched bases divided by read length) over the reads not yet covered. Segments of one genome recruit different reads and are all selected. Strains of the same segment share reads, so only the best is selected. A primary pick is replaced by the overlapping secondary pick only if the latter aligns better on the shared reads by more than one percentage point. Secondary picks that do not overlap any primary pick are added. The selection and the reason for every candidate’s inclusion or exclusion are written to ref_selection.tsv.

### Consensus assembly

All host-filtered reads are mapped to the concatenated references with minimap2 (-k 11 -w 5). Alignments are sorted, fixmate-corrected and duplicate-marked with SAMtools^34^. Positions with depth below min_coverage (from samtools depth -J, which counts deletions as covered) are written to a mask. Variants are called with bcftools mpileup -X ont-sup -B, bcftools call --ploidy 1 -mv and bcftools norm -m -any. A variant table lists alleles with allele frequency ≥ report_min_af and ≥ min_ac supporting reads. Only alleles with frequency > 0.5 are applied to the consensus by bcftools consensus, and masked positions become N. min_coverage defaults to 10, used by all benchmarks except where stated.

### Consensus polishing

For ONT reads the consensus is then polished with Medaka^29^ (v2.2.2). A draft is built by applying the same dominant alleles to the unmasked reference, because reads cannot align across masked stretches. The reads used for the consensus are mapped to the draft with the same minimap2 settings. medaka inference is run on the draft regions with depth ≥ min_coverage, and medaka sequence writes the polished sequence with all other positions set to N. The reads are remapped to the polished sequence and positions below min_coverage are masked again. The Medaka model comes from the read headers, or from the configuration when they carry none.

### Real-time mode

emtrace watch monitors a directory with the watchdog library. It enqueues each new FASTQ file after its size has been stable for 2 s, because MinKNOW writes files incrementally. It then runs preprocessing, host filtering, annotation and EM on each file, merges the annotation with earlier batches and rewrites the cumulative abundance table. A final report is written when monitoring stops.

### Interfaces and distribution

The command-line (emtrace run), graphical (emtrace gui, PyQt6) and real-time (emtrace watch) interfaces share one configuration schema. Databases are downloaded once with a built-in manager. EMtrace is released under the GNU GPLv3 and is installed either from source with pip, with its external tools from Bioconda, or as a self- contained macOS desktop bundle attached to releases from v1.0.3 onward (Code availability).

### Benchmark design and scoring

The benchmark is organized into frameworks, each with its own question, datasets, truth and comparators (Supplementary Table 1). All scoring used ngs-benchmark v1.0.0, a Python library developed for this study and released with the benchmark (Code availability). It reads each tool’s native output through an adapter, so every tool is scored by the same code.

### Detection (framework A)

Detection was scored at species rank, per sample, against per- sample truth. A taxon was called when it had at least 10 reads, counted with each tool’s own estimate: readn_em for EMtrace, Bracken’s new_est_reads for EPI2ME, the larger of CZ ID’s nucleotide- and protein-level counts (nt_count, nr_count), and the read count of MEGAN-LR’s taxon export (read-count mode). A taxon also had to hold at least 0.01% of the sample’s classified non-host reads, the abundance floor used for 16S profiling and by Emu^14^. Host reads (division Primates) were left out of this total, because EPI2ME reports human reads as taxa, whereas EMtrace and CZ ID remove them before classification. True positives, false positives and false negatives were pooled over the samples of a dataset before computing precision, recall and F1 (micro-averaging). EMtrace’s calls on the field libraries and clinical samples were taken from its framework B runs with the default annotation, which use the same configuration and host filter.

### Assembly (framework B)

Each consensus was aligned to the scoring genome with MAFFT^35^ (--globalpair --maxiterate 1000 --adjustdirection). Coverage is the percentage of scoring-genome positions aligned to a non-N consensus base. Identity is the percentage of identical bases over aligned non-gap, non-N columns. Substitution errors are mismatches over the same columns. Indels are counted from gaps in consensus and reference sequences, each indel counted once whatever its length. A sample without a consensus scores 0% coverage and is excluded from identity and error rates. For the simulated divergent genomes, the true genome is the genome from which the reads were simulated. For the field libraries, where the true genome is unknown, it is the closest complete genome to the sequenced isolate (Supplementary Table 5). For the 16 SARS-CoV-2 clinical samples, which span several lineages and have no isolate genome in GenBank, it is Wuhan-Hu-1 (NC_045512.2). Their identity therefore includes lineage differences from this reference. Where EMtrace assembled two accessions of one virus, the record with the most reads was scored.

### Real-time replay (framework C)

Each library was replayed as 4,000-read files (MinKNOW’s default) in order of read start time; the tick pools, whose SRA reads carry no start times, were cut in file order and scored without a sequencing-time axis. Each batch file was released into the watched directory once the previous file’s abundance table had been written, and each table was scored with ngs-benchmark’s realtime module. For each batch, sequencing time is its latest read start_time minus the run’s earliest, and latency the wall-clock time from file release to the updated table. Time to first correct call is the first batch from which the top-ranked taxon stays correct. To reconcile the original study’s virus names with NCBI taxonomy, which classifies Aigai virus separately from CCHFV, the tick pools were scored at genus rank against a panel of that study’s findings. A PCR-positive virus counts as detected, and a PCR-negative one as a false detection, once it reaches 10 reads and stays at or above that count for the rest of the run. Viruses that the study reported from nanopore sequencing alone are scored as found or not found; precision and recall are not reported for them. Stabilisation is the first batch after which L1 stays within ±0.01 of its final value. Per-batch precision and recall use the framework A definitions. Genome coverage over time was computed after the replays from their per-batch host-filtered reads: the reads of batches 1…*k* were pooled at log-spaced *k* (up to 20 points; all 12 batches of pool 58-37) and assembled with --min_coverage 10 and no polishing against KEVOOV fi663 (PQ165789–PQ165798), TBEV Neudoerfl (NC_001672.1), CCHFV (Aigai virus) isolate CAP12 (MN811031, MN811037, MN811034) and JMTV strain Tulcea1 (MW561147– MW561150). CAP12 and Tulcea1 are complete coding sequences, the nearest complete relatives in the original study’s phylogenies. The final KEVOOV and TBEV assemblies were also polished with Medaka; the tick pools, whose basecalling model is not recorded, were not. Each segment was scored as in framework B, and whole-genome coverage is length-weighted over the segments.

### 16S profiling

Abundances were scored against the 16S-copy-number-corrected truth of each mock community, with near-synonyms collapsed by a per-library map (EMtrace-benchmark repository) and a 10-read threshold on each tool’s own estimate. Uncollapsed scores are reported alongside. L1 is the sum of absolute differences between predicted and true relative abundances over the union of taxa. Predicted abundances are each tool’s own estimate: the EM estimate *F*(*t*) for EMtrace and Emu’s EM estimate, and read counts for Kraken 2, Bracken and Spaghetti. *Other* is the percentage of reads assigned to taxa outside the truth. After the 10-read threshold, taxa below 0.01% relative abundance were removed for every tool, the floor Emu’s authors applied to all methods on the ZymoBIOMICS standard^14^ (Table 1, Supplementary Table 4).

## Datasets

### Field libraries

Four ONT libraries used in the ViMOP benchmark^6^: Zika virus (SRR31174237), West Nile virus (ERR14941561), SARS-CoV-2 (SRR15356294) and dengue virus 2 (SRR32419728).

### SARS-CoV-2 clinical samples

Sixteen nasopharyngeal samples with RT-qPCR Ct values from BioProject PRJNA761346 (ref. 25). Five negative controls were run but not scored.

### Viral mock community

The ATCC MSA-1008 mock in human background from Buddle et al.^11^, untargeted ONT DNA and RNA libraries at 60, 600, 6,000 and 60,000 genome copies per mL. One library per concentration and nucleic-acid type was used (RNA ERR13488556, ERR13488558, ERR13488560 and ERR13488562; DNA ERR13485822, ERR13488550, ERR13488553 and ERR13488749).

### Simulated divergent viruses

Divergent genomes and reads were generated following the design of the ViMOP benchmark^6^, with our own implementation (scripts in the EMtrace- benchmark repository). The source genomes were SARS-CoV-2 OX637002.1, dengue virus 2 NC_001474.2 and Lassa virus L and S (MG812630.1, MG812631.1). For each divergence level r ∈ {0, 5, 10, 15, 20, 25, 30}%, ceil(*r* × *L*) positions of a source genome of length L were drawn without replacement. Draws were weighted by the number of sequences that differ from the column consensus in ViMOP’s curated alignment of that virus, and each drawn base was replaced by one of the other three bases chosen uniformly (seed 42). Reads were simulated with PBSIM3^36^ in sample-based mode, using ViMOP’s SISPA read profile, at 30× mean depth, and given MinION-style headers. Background reads were simulated in the same way from the human genome (GRCh38.p14) and eight respiratory bacteria; 500,000 of them, in ViMOP’s proportions (75% human), were added to each viral read set without downsampling it.

### Replay libraries

KEVOOV isolate fi663, from *Ochlerotatus communis* mosquitoes collected in Finnish Lapland and isolated in C6/36 cells^32^: dsRNA amplified by full-length amplification of cDNA (FLAC) and sequenced on a MinION Mk1C (R9.4.1, SQK-LSK110; model 2021-05- 17_dna_r9.4.1_minion_96; 464,000 reads replayed over 19.0 h). TBEV: four *Ixodes ricinus* tick samples (IL209-11-15, IL232-5, IL239-1, IL256-1) collected in the Helsinki area, Finland, in 2025: random-hexamer cDNA amplified by TBEV-specific tiled-amplicon PCR^37^, barcoded and sequenced together on a MinION Mk1D (R10.4.1, SQK-RBK004; model dna_r10.4.1_e8.2_400bps_hac@v5.2.0; 313,867 reads replayed over 5.0 h). KEVOOV and TBEV are each a single contiguous run. Tick pools 58-37 and 35-33 (BioProject PRJNA878809; SRR21504617, 44,503 reads, and SRR21504592, 216,828 reads): seven female *Rhipicephalus bursa* each, random-primed cDNA without amplification, GridION. Both reported CCHFV PCR- positive; 58-37 is also JMTV PCR-positive and 35-33 JMTV PCR-negative^33^. The 16S library barcode79 (below; 26 files, 71.8 h) was also replayed.

**16S mock communities.** barcode79, a ZymoBIOMICS microbial community standard (R6300, eight bacterial species; flow cell FAS57016; R9.4.1 basecalled with Guppy (super-accuracy model); 100,128 reads), and the 21-species synthetic gut community of Curry et al.^14^ (SRR14326315; 521,661 reads). The published ONT abundance column was used as truth. The ZymoBIOMICS Gut Microbiome Standard (D6331; 14 bacterial species, including five *Escherichia coli* strains, one archaeon and two yeasts) came from ONT’s open dataset zymo_16s_2025.09^30^ (s3://ont-open-data/zymo_16s_2025.09): four SQK-MAB114.24 libraries, two biological replicates each sequenced on two R10.4.1 flow cells, basecalled by ONT with Dorado 1.1.1 (SUP model v5.2.0; 72,830–84,233 reads). The certificate’s 16S-only composition was used as truth, with the five *E. coli* strains summed and the yeasts at 0%.

### Comparator configuration

**EPI2ME wf-metagenomics** v2.14.2 (Kraken 2 + Bracken, with the workflow’s Standard-8 database set: the Kraken 2 Standard index capped at 8 GB, release 2024-12-28, built from RefSeq archaea, bacteria, viruses, plasmids and human plus UniVec_Core, with NCBI taxonomy of 2025-01-01), **CZ ID** (replaced by SeqtoID since October 2026; public cloud, mNGS Nanopore pipeline, samples processed 2026-08-06) and **MEGAN-LR** (MEGAN6 Community Edition v6.25.10, long-read mode; LAST v1652 frameshift-aware DNA-to-protein alignment with a custom protein database: RefSeq release 234, built from all viral, one archaeal/bacterial genome per genus and human proteins, with accessions mapped to NCBI taxonomy via megan- map-Jul2020-2; human reads removed beforehand with nf-core/taxprofiler v2.0.1 (minimap2 against GRCh38.p14)), were run with default settings. **ViMOP** (v1.0.5, commit 74dae412ec) was run with Apptainer (Nextflow 25.10.2) on CSC’s Puhti supercluster and, for the field libraries, in Docker (Nextflow 26.04.3) on the laptop, with its own reference set, and scored on its own consensus. ViMOP’s benchmark settings (0 nt read-end trim, 20× minimum coverage, 50% minimum genome recovery) cannot be set there. **INSaFLU-TELEVIR** 2.2.2 was run on its public web server (Classic workflows; accessed 28 August – 2 September 2026) with every pipeline stage at the instance defaults (per-stage tools, versions and parameters in Supplementary Table 6). TELEVIR returns a read alignment without a depth-thresholded consensus, so its consensus was built from its alignment with samtools consensus -m simple -d 10 -c 0.5, which matches EMtrace’s 10× threshold and majority rule. **Emu** (v3.6.2) was run with its own database at default settings. **Spaghetti** (commit ac9d2ec; Porechop 0.2.4, NanoFilt 2.8.0 with Spaghetti’s 1,200–1,800 bp length window) was run on SILVA 138 (QIIME 2 release) with its chimera-removal step (yacrd on an all-versus-all minimap2 alignment) skipped. The size of this alignment grows with the square of the read count: scaled from a 5,000-read subsample, about 490 GB for the ZymoBIOMICS mock and 13.8 TB for the synthetic gut community, beyond the storage of most portable computers. Neither Emu nor EMtrace removes chimeras, so skipping the step also keeps the comparison like for like.

**Kraken 2** and **Bracken** were run on Kraken 2’s Greengenes 16S build. wf-16s (EPI2ME, ONT) was run on the R10.4.1 gut community only, with Nextflow and Docker, its minimap2 classifier and default ncbi_16s_18s database; species abundances were counted from its per-read assignments, because the release used reports at genus rank by default. For the shared- reference comparison, the 49,243 16S sequences of Emu’s default database were indexed unchanged for EMtrace, Spaghetti and Kraken 2/Bracken. Each tool applied its own read preprocessing. EMtrace’s Q10 filter kept 62% of the synthetic-gut reads, whereas Emu and Spaghetti applied no quality filter.

### EMtrace configurations

**Virome:** host filter *Homo sapiens* only (frameworks A and B; the replays below added the kitome). **Default annotation:** minimap2 against minimap.refseq.abv.1g (RefSeq release 234 archaea, bacteria and viruses, one archaeal/bacterial genome per species; *k* = 15, *w* = 10).

**Chain2:** the same, then DIAMOND BLASTP against diamond.uniref.vi (UniRef100 release 2026_01, viral proteins) on orfipy ORFs ≥90 nt of the reads left unannotated, bitscore ≥80.

Reference selection for chain2 used the primary source only (--no-ref_auto_annotation). Viral replays: minimap.refseq.abv.1g with k = 11, w = 5 (prebuilt index) for KEVOOV and the tick pools, alone and followed by DIAMOND as in chain2; the default index for TBEV. Host filter kitome plus *H. sapiens* and the host of each library (*Aedes albopictus* for KEVOOV, *Ixodes ricinus* and *I. persulcatus* for TBEV, *Rhipicephalus sanguineus* for the tick pools, the nearest relative of *R. bursa* with a genome assembly). Porechop_ABI, Chopper Q8 and ≥150 bp (KEVOOV) or ≥100 bp without read splitting (TBEV and tick pools). **Consensus:** min_coverage 10, Medaka polishing. **16S:** Porechop_ABI, Chopper Q10 and 1,200–1,800 bp, minimap2 against de-duplicated rrnDB (16S.rrnDB.dedup) or Emu’s reference, -N 25, all hits, EM temperature 10. Defaults were chosen in preliminary tests and used unchanged throughout.

Configuration files for every run are provided in the EMtrace-benchmark repository (jobs/).

### Hardware

Framework A analyses ran on the CSC Roihu HPC (2 × 192-core AMD EPYC 9965, Zen 5, 768 GiB RAM per node, Finland), except EMtrace on the field libraries and clinical samples, whose framework B laptop runs were reused (Detection, above). Framework B ran on an Apple M2 Pro laptop (12 cores, 16 GB RAM) and on CSC’s Puhti, a Linux/Unix CPU supercluster with 24 cores per ViMOP run, each running at 2.1 GHz. The framework C replays and all 16S tools (EMtrace, Emu, Spaghetti and Kraken 2) were run and timed on the laptop with 8 threads.

Runtime and peak memory per tool and dataset are given in Supplementary Table 7; values compare tools only within a hardware class.

### Statistics and reproducibility

Datasets were small (4–16 samples per detection dataset, one library per condition). Results are reported descriptively, without hypothesis tests. No data were excluded except the negative controls of the clinical SARS-CoV-2 study, which have no viral truth.

## Data availability

Sequencing data are available from SRA/ENA under the accessions listed in Methods. The KEVOOV and TBEV replay libraries are available from the authors on request. The simulated divergent genomes and reads were generated for this study following the design of the ViMOP benchmark^6^ (Methods) and are available, with the scripts that generated them, at Zenodo (https://doi.org/10.5281/zenodo.23978393). Truth tables, scoring references, per-sample metrics and figure source data are in the EMtrace-benchmark repository (Code availability).

## Code availability

The EMtrace source code is freely available at https://github.com/weinstein-bioinfo/EMtrace under the GNU General Public License v3.0 or later. The version used in this study, v1.0.0, is archived at Zenodo (https://doi.org/10.5281/zenodo.24079771). Later releases, including updates to the desktop interface and macOS bundles, are available from the repository. All code needed to reproduce the benchmark is freely available at https://github.com/weinstein-bioinfo/EMtrace-benchmark under the same license and is archived at Zenodo (https://doi.org/10.5281/zenodo.23976647).

## Acknowledgements

The authors wish to acknowledge the CSC IT Center for Science, Finland, for providing computational resources.

## Author contributions

Conceptualization of the project: I.W., T.Sm. and R.K.; Conceptualization of the pipeline: I.W., T.Sm. and R.K.; Implementation, updates, and code distribution: I.W.; Reference database curation: I.W.; Investigation (sequencing data): M.K. and H.V.; Benchmarking: I.W., O.L. and M.K.; Testing and reporting: I.W., O.L., M.K., T.Sm. and R.K.; Writing—original draft preparation: I.W.; Writing—review and editing: O.V., R.K., T.Sm. and M.K.; Supervision of the study: R.K., T.Sm. and O.V.; Funding acquisition: R.K., T.Si., M.G. and O.V.

