## Supplementary Information: Supplementary Tables 1-7, Supplementary Figure 1 and Supplementary Note 1 for "EMtrace: offline nanopore metagenomics on a laptop for rapid viral diagnostics, genome recovery and 16S profiling"

### Contents

| | |
|---|---|
| Supplementary Table 1 | Benchmark design: datasets, tools and the framework each pairing was scored under |
| Supplementary Table 2 | Sub-threshold and per-tool read counts on the viral mock community |
| Supplementary Table 3 | Genome recovery and consensus accuracy for simulated divergent viruses |
| Supplementary Table 4 | 16S profiling with each tool on its own database |
| Supplementary Table 5 | Scoring references for the real libraries |
| Supplementary Table 6 | INSaFLU-TELEVIR configuration |
| Supplementary Table 7 | Runtime and memory per tool and dataset |
| Supplementary Figure 1 | Genome recovery from real libraries |
| Supplementary Note 1 | Host-filter alignment-score threshold on simulated reads |

### Supplementary Table 1 | Benchmark design

**A, B, C** mark the framework each dataset–tool pairing was scored under: **A** detection accuracy, **B** consensus assembly, **C** real-time replay. **16S** marks the 16S profiling track. A dash means the tool was not run on that dataset. EMtrace was run on every dataset; no comparator analyses reads during sequencing, so framework C has no comparator column entries.

| Dataset (samples scored) | EMtrace | EPI2ME wf-metagenomics | CZ ID | MEGAN-LR | metaMix | ViMOP | INSaFLU-TELEVIR | Emu | Spaghetti | Kraken 2/ Bracken | EPI2ME wf-16s |
|---|---|---|---|---|---|---|---|---|---|---|---|
| **Virome** | | | | | | | | | | | |
| Field libraries, Petersen et al. (4) | A, B | A | A | A | A* | B | B | – | – | – | – |
| SARS-CoV-2 clinical, Gauthier et al. (16) | A, B | A | A | A | A* | B | B | – | – | – | – |
| Viral mock community, Buddle et al. (8) | A | A | A | A | A* | – | – | – | – | – | – |
| Simulated divergent viruses, pure (28) | B | – | – | – | – | B | B | – | – | – | – |
| Simulated divergent viruses, mixed (28) | B | – | – | – | – | B | B | – | – | – | – |
| **Real-time replay** | | | | | | | | | | | |
| KEVOOV, in-house (1) | C | – | – | – | – | – | – | – | – | – | – |
| TBEV, in-house (1) | C | – | – | – | – | – | – | – | – | – | – |
| Tick pools, Ergunay et al. (2) | C | – | – | – | – | – | – | – | – | – | – |
| **16S rRNA** | | | | | | | | | | | |
| Synthetic gut community, Curry et al. (1) | 16S | – | – | – | – | – | – | 16S | 16S | 16S | – |
| ZymoBIOMICS mock, barcode79 (1) | 16S, C | – | – | – | – | – | – | 16S | 16S | 16S | – |
| ZymoBIOMICS Gut Microbiome Standard (4) | 16S, C | – | – | – | – | – | – | 16S | 16S | 16S | 16S |

**Supplementary Table 1 | Benchmark design: datasets, tools and the framework each pairing was scored under.** The benchmark is split into frameworks because the four questions need different data, truth and comparators. **Framework A**, detection accuracy: species-rank precision, recall and F1 against per-sample truth, pooled over each dataset's samples. **Framework B**, consensus assembly: coverage, identity, and substitutions and indels per 10 kbp against a fixed scoring genome per sample. **Framework C**, real-time replay: time to first correct call, per-batch compute time and throughput ratio, and genome coverage or abundance error over the run; no comparator processes reads during sequencing (Figure 1c), so EMtrace is compared against the run length a post-run tool would have to wait for. **16S profiling**: genus- and species-rank L1 error, false positives and false negatives, with every tool on one shared reference (Table 1) and on its own database. Dataset accessions, library details and scoring definitions are in Methods. *metaMix was run on the three detection datasets but made no species-level call on any sample, so it is not scored.

### Supplementary Table 2 | Sub-threshold and per-tool read counts on the viral mock community

#### a. Reads per spiked virus at 60,000 genome copies per mL, the only load at which any tool crossed the 10-read threshold.

| Library | Virus (spiked) | EMtrace | EPI2ME | CZ ID (nt / nr) | MEGAN-LR |
|---|---|---|---|---|---|
| RNA | *Orthoflavivirus zikaense* | 45 | 45 | 45 / — | 9 |
| RNA | *Orthopneumovirus hominis* | 24 | 26 | 25 / 5 | 5 |
| RNA | *Orthoreovirus mammalis* | 7 | — | 6 / — | 3 |
| RNA | *Betainfluenzavirus influenzae* | 1 | — | 1 / — | — |
| DNA | *Cytomegalovirus humanbeta5* | 96 | 98 | 1 / 75 | 17 |
| DNA | *Mastadenovirus faecale* | 44 | 44 | 44 / 37 | 42 |

#### b. Taxa EMtrace reported below the 10-read threshold, at every load.

| Library | Load (gc/mL) | Sample | Taxon | Reads | Spiked |
|---|---|---|---|---|---|
| RNA | 6,000 | ERR13488558 | *Orthopneumovirus hominis* | 9 | yes |
| RNA | 6,000 | ERR13488558 | *Orthoflavivirus zikaense* | 2 | yes |
| DNA | 6,000 | ERR13488550 | *Cytomegalovirus humanbeta5* | 5 | yes |
| DNA | 600 | ERR13488749 | *Cytomegalovirus humanbeta5* | 5 | yes |
| RNA | 60 / 600 / 6,000 | ERR13488562 / 560 / 558 | BeAn 58058 virus | 1 / 3 / 10 | no |
| DNA | 60 | ERR13488553 | BeAn 58058 virus | 3 | no |
| DNA | 600 | ERR13488749 | *Simplexvirus atelinealpha1* | 1 | no |

#### c. Libraries by viral load.

| Load (gc/mL) | RNA library | DNA library |
|---|---|---|
| 60,000 | ERR13488556 | ERR13485822 |
| 6,000 | ERR13488558 | ERR13488550 |
| 600 | ERR13488560 | ERR13488749 |
| 60 | ERR13488562 | ERR13488553 |

**Supplementary Table 2 | Sub-threshold and per-tool read counts on the viral mock community.** The ATCC MSA-1008 six-virus mock in human background, untargeted ONT DNA and RNA libraries at four viral loads, one library per load and nucleic-acid type (Methods). Counts are each tool's own raw output before the 10-read threshold and 0.01% abundance floor were applied: EMtrace `readn_em`, Bracken `new_est_reads` for EPI2ME, CZ ID `nt_count` / `nr_count`, and the read-count export for MEGAN-LR. An em dash means the taxon was absent from that tool's output. **a**, two of the six spiked viruses fall below the threshold even at the highest load (*Orthoreovirus mammalis*, 7 reads; *Betainfluenzavirus influenzae*, 1 read), so with the sub-threshold rows of **b** EMtrace recovered spiked viruses at the three highest loads. **b**, BeAn 58058 virus and *Simplexvirus atelinealpha1* are not part of the mock; neither reaches the threshold except BeAn 58058 virus at 6,000 gc/mL in the RNA library. Detection in this mock is therefore limited by the number of viral reads rather than by the classifier.

### Supplementary Table 3 | Genome recovery and consensus accuracy for simulated divergent viruses

| Tool | 0% | 5% | 10% | 15% | 20% | 25% | 30% | Mean, pure | Mean, mixed | Subst/10 kbp | Indels/10 kbp |
|---|---|---|---|---|---|---|---|---|---|---|---|
| EMtrace | **92.9** | **92.5** | 70.2 | 60.0 | 57.0 | 42.3 | 27.3 | 63.2 | 63.2 | **3.6** | **1.9** |
| EMtrace (chain2) | **92.9** | **92.5** | **92.4** | **79.7** | **72.5** | **57.3** | **36.6** | **74.9** | **74.9** | **3.6** | **1.9** |
| ViMOP | 84.2 | 84.0 | 83.3 | 73.1 | 61.0 | 36.8 | 27.0 | 64.2 | 64.2 | 5.3 | 3.7 |
| INSaFLU-TELEVIR | 89.1 | 71.5 | 48.5 | 50.8 | 46.6 | 32.9 | 30.3 | 52.8 | 40.5 | 8.6 | 3.9 |

**Supplementary Table 3 | Genome recovery and consensus accuracy for simulated divergent viruses.** Genome coverage (%) is the mean over four target sequences per divergence level, pure reads. Mean columns average all 28 pure or mixed samples, counting a sample with no consensus as 0%. Substitutions and indels per 10 kbp of reported bases are pooled over the pure samples with a consensus. Identity over the reported bases was ≥99.9% for every tool. EMtrace consensus polished with Medaka. Bold, best per column.

### Supplementary Table 4 | 16S profiling with each tool on its own database

| Library | Tool | Database | Genus L1 | FP genera | FN genera | Species L1 | FP species | FN species | Species precision |
|---|---|---|---|---|---|---|---|---|---|
| Synthetic gut | Kraken 2 | Greengenes 16S | 1.254 | 55 | 9 | 1.413 | 43 | 17 | 0.07 |
| Synthetic gut | Bracken | Greengenes 16S | 1.449 | 28 | 11 | 1.413 | 58 | 17 | 0.05 |
| Synthetic gut | Spaghetti | SILVA 138 | 0.349 | 26 | **1** | — | — | — | — |
| Synthetic gut | Emu | Emu (49,243 seqs) | **0.149** | 10 | **1** | 0.500 | 26 | **3** | 0.40 |
| Synthetic gut | EMtrace | rrnDB 5.10, de-duplicated | 0.351 | **4** | **1** | **0.446** | **12** | **3** | **0.59** |
| ZymoBIOMICS mock | Kraken 2 | Greengenes 16S | 1.051 | 37 | 3 | 1.571 | 69 | 6 | 0.03 |
| ZymoBIOMICS mock | Bracken | Greengenes 16S | 1.143 | 14 | 3 | 1.538 | 67 | 6 | 0.03 |
| ZymoBIOMICS mock | Spaghetti | SILVA 138 | 0.291 | 13 | **0** | — | — | — | — |
| ZymoBIOMICS mock | Emu | Emu (49,243 seqs) | **0.254** | **10** | **0** | **0.254** | 28 | **0** | 0.22 |
| ZymoBIOMICS mock | EMtrace | rrnDB 5.10, de-duplicated | 0.259 | **10** | **0** | 0.255 | **12** | **0** | **0.40** |
| Gut Microbiome Standard | Kraken 2 | Greengenes 16S | 0.870 | 26.8 | 5.0 | 1.401 | 36.3 | 9.0 | 0.12 |
| Gut Microbiome Standard | Bracken | Greengenes 16S | 0.958 | 9.8 | 6.0 | 1.391 | 34.3 | 9.0 | 0.13 |
| Gut Microbiome Standard | Spaghetti | SILVA 138 | 0.586 | 4.0 | **2.5** | — | — | — | — |
| Gut Microbiome Standard | Emu | Emu (49,243 seqs) | 0.585 | 1.0 | **2.5** | 0.873 | 5.8 | **2.5** | 0.67 |
| Gut Microbiome Standard | wf-16s (ONT) | NCBI 16S/18S | 0.584 | **0.0** | 2.8 | 1.359 | 11.3 | 3.8 | 0.48 |
| Gut Microbiome Standard | EMtrace | rrnDB 5.10, de-duplicated | **0.581** | **0.0** | 2.8 | **0.581** | **0.0** | 2.8 | **1.00** |

**Supplementary Table 4 | 16S profiling with each tool on its own database.** Synthetic gut community (21 species; 521,661 reads), ZymoBIOMICS mock (8 species; 100,128 reads) and ZymoBIOMICS Gut Microbiome Standard (14 bacterial species; R10.4.1; mean over four replicate libraries). Each tool with its default database, synonyms collapsed, 10-read threshold and taxa below 0.01% relative abundance removed for every tool, as in Table 1 (on the ZymoBIOMICS mock the floor removes nothing beyond the 10-read threshold). EMtrace in its frozen configuration (-N 25, EM temperature 10), as in Table 1. Spaghetti reports genera only. wf-16s, ONT's workflow with its minimap2 classifier (Gut Microbiome Standard only), reports through an NCBI taxonomy from 2024 that splits *Faecalibacterium prausnitzii* into species described since 2022 and names *Fusobacterium nucleatum* reads *F. animalis*; these count as false positives against the certified names. Without the floor, Emu reported 52 false-positive species on the synthetic gut and EMtrace 13. FP, false positives; FN, false negatives. Bold, best per column within each library.

### Supplementary Table 5 | Scoring references for the real libraries

| Library | Virus | Isolate | Isolate genome in GenBank | Scoring reference | Length (bp) | Identity to isolate genome |
|---|---|---|---|---|---|---|
| SRR31174237 | Zika virus | R116265, Mexico 2016, Asian lineage, C6/36 lab stock | KX766029.1, lacks both genome ends | **MT507047.1**, S-542, Yucatán 2017 | 10,807 | 99.77% |
| SRR32419728 | Dengue virus 2 | R0908/08, Thailand 2008, genotype Asian I | PQ810041.1, complete CDS only | **DQ181797.1**, ThD2_0078_01, Thailand 2001 | 10,723 | 99.23% |
| SRR15356294 | SARS-CoV-2 | Oxford, April 2020 (Ct 29.9) | none | **NC_045512.2**, Wuhan-Hu-1 | 29,903 | — |
| ERR14941561 | West Nile virus | strain B956, lineage II, lab stock | none | **NC_001563.2**, strain 956 | 10,962 | — |
| PRJNA761346 (16) | SARS-CoV-2 | 16 nasopharyngeal samples, several lineages | none | **NC_045512.2**, Wuhan-Hu-1 | 29,903 | — |

**Supplementary Table 5 | Scoring references for the real libraries.** The **scoring reference** is the single genome that every pipeline's consensus is aligned to when coverage and identity are measured (Methods). A real library has no known true genome, so one fixed reference per sample is chosen in advance and independently of any pipeline, and the same one is used for EMtrace, ViMOP and INSaFLU-TELEVIR alike. Two rules decide it: the reference must be a **complete** genome, with both ends present, and among complete genomes the one closest to the sequenced isolate is used. **Isolate genome in GenBank** is a published genome of the same isolate, identified from each library's BioSample metadata. Both are partial, so neither could serve as the scoring reference. **Identity to isolate genome** is the scoring reference's percentage identity to that genome, over aligned non-gap columns; candidates came from BLAST (megablast, core_nt, full-length genomes only). Bold, the reference used for scoring. Where an isolate genome exists, the scoring reference is 99.2–99.8% identical to it, so identity measured against the scoring reference approximates consensus accuracy.

**Rationale per library.**

- **Zika.** The BioSample's `/isolate` field gives the accession KX766029.1, but that record lacks both genome ends (10,749 bp) and is therefore not used. MT507047.1 is a Mexican isolate of the same lineage; four slightly closer records (up to 99.81%) are truncated at one end, and MT507048.1 is a clone of the same isolate, so the parent isolate was taken. The RefSeq genome NC_012532.1 (MR766, Uganda) is African lineage and only 88.9% identical to the isolate.
- **Dengue virus 2.** PQ810041.1, a GenBank record linked to the same BioSample, is filed as complete CDS with a partial 3′ region and is therefore not used. DQ181797.1 is the closest full-length genome, Thai and of the same genotype.
- **SARS-CoV-2 field library.** No genome of this sample is in GenBank. The library is a UK sample from April 2020, close to the ancestral lineage, so Wuhan-Hu-1 is used.
- **West Nile virus.** The library is the B956 strain and RefSeq NC_001563.2 is that strain. All three pipelines score 99.7% identity against it, so the lab stock may carry passage differences from the RefSeq sequence; with no isolate genome to decide, RefSeq is kept.
- **SARS-CoV-2 clinical samples.** The 16 samples span several lineages (2020 early lineages, 2021 presumptive Alpha and N501Y + E484K variants) and none has an isolate genome in GenBank, so all are scored against Wuhan-Hu-1. Their identity therefore includes lineage differences from this reference.

### Supplementary Table 6 | INSaFLU-TELEVIR configuration

INSaFLU-TELEVIR was run through its free public web instance (https://insaflu.insa.pt), the interface through which the tool is delivered. All 76 samples of the assembly benchmark (28 pure and 28 background-embedded simulated mutants, 4 field libraries, 16 SARS-CoV-2 clinical samples) were uploaded as ONT FASTQ files and run with the settings below. For the most divergent Lassa virus samples the assembly step returned no contigs, so those runs produced no consensus to download and score 0% genome coverage, as for any sample without a consensus (Methods).

| Field | Value |
|---|---|
| Instance | INSaFLU public web server, https://insaflu.insa.pt |
| INSaFLU version | 2.2.2 |
| Access date | 28 August 2026 |
| Workflow | Run → Classic workflows |
| Parameters | Every pipeline stage at the public-instance defaults (below); no other settings are exposed by the interface |
| Host-depletion reference | Human GRCh38.p14 (`GCF_000001405.40_GRCh38.p14_genomic.fna.gz`) |
| Classification databases | Centrifuge `viral_index`; BLASTN `refseq`; DIAMOND `refseq_prot`; Kaiju `kaiju_db_viruses.fmi`; KrakenUniq `viral`. Database build dates are not reported by the interface. |

**Pipeline stages, tool versions and parameters**, as deployed on the public instance.

| Stage | Tool (version) | Parameters | Used |
|---|---|---|---|
| Extra QC (low complexity) | Prinseq++ v1.2.4 | `--lc_entropy 0.5 --lc_dust 0.7` | yes |
| Viral enrichment | Centrifuge v1.0.4_beta | `--min-hitlen 20 -k 5 --db viral_index --min-hits 1` | yes |
| Host depletion | Minimap2 v2.24 | `-x map-ont --secondary=no`, against GRCh38.p14 | yes |
| Assembly | Raven v1.8.1 | `-f 0.001 -w 5 -k 21 -p 2` | yes |
| Contig classification | BLASTN v2.12.0 | `-max_target_seqs 5 -evalue 1e-5 --db refseq` | yes |
| Contig classification | Kraken2 v2.1.2 | `--quick ON --confidence 0.5 --db viral --min-hits 1` | no (alternative) |
| Read classification | DIAMOND v2.0.15 | `--fast --id 40 --query-cover 40 --db refseq_prot` | yes |
| Read classification | Kaiju v1.8.2 | `-e 3 -s 65 -X --db kaiju_db_viruses.fmi` | yes |
| Read classification | KrakenUniq v0.7.3 | `--db viral` | yes |
| Remapping | Minimap2 v2.24 | `-ax map-ont --secondary=no` | yes |
| Remap filtering | — | none (BamUtil, dustmasker and msamtools available, not selected) | no |
| Validation, request mapping | Minimap2 v2.24 | `-ax map-ont --secondary=no` | yes |
| Validation, screening | Minimap2 v2.24 | `-ax map-ont --secondary=no` | yes |
| Reporting, flagging and sorting | Final Report v1.0.0 | `--flag-type viruses --r-overlap 0.5` | yes |
| Reporting, remap management | v1.0.0 | `--max-taxids 12 --max-accids 4 --include-manual ON` | yes |

**Settings not available on the public instance.** The ViMOP benchmark[6] ran TELEVIR on a local installation with three settings changed from their defaults: read-end trimming 0 nt (default 30 nt), minimum consensus coverage 20×, and minimum genome recovery 50%. None of these can be set through the public web interface, whose only exposed controls are the Prinseq++ complexity filter and the Raven assembly parameters above. Our runs therefore use the platform's fixed values for all three.

**Consensus used for scoring.** TELEVIR returns, per detected virus, the reference it mapped to and the mapped-read alignment (BAM), without a depth-thresholded consensus sequence. For scoring, the consensus was built from that alignment with `samtools consensus -m simple -d 10 -c 0.5`: positions below 10× depth are N, and a base is called when it is the majority allele (>50%). This matches the 10× threshold and majority rule applied to EMtrace. Samples for which TELEVIR returned no alignment for the target virus count as 0% genome coverage.

### Supplementary Table 7 | Runtime and memory per tool and dataset

| Dataset (samples) | Tool | Hardware (threads) | Wall time per sample, median (range) | CPU-hours per sample, median | Peak RAM, max (GB) |
|---|---|---|---|---|---|
| **Virome** | | | | | |
| Simulated mutants, pure (28) | EMtrace | Laptop (8) | 1.3 min (0.6–1.6) | 0.02 | 6.9 |
| | EMtrace (chain2) | Laptop (8) | 1.7 min (1.5–2.2) | 0.06 | 7.3 |
| | ViMOP | Puhti (24) | 8.2 min (0.5–34.0) ‡ | 0.18 | 12.2 |
| | INSaFLU-TELEVIR | public web server | n.m. | n.m. | n.m. |
| Simulated mutants, mixed (28) | EMtrace | Laptop (8) | 9.1 min (7.9–11.7) | 0.47 | 9.5 |
| | EMtrace (chain2) | Laptop (8) | 13.6 min (11.8–14.8) | 1.04 | 9.5 |
| | ViMOP | Puhti (24) | 5.0 h (4.5–5.7) ‡ | 14.2 | 21.0 |
| | INSaFLU-TELEVIR | public web server | n.m. | n.m. | n.m. |
| Field libraries, Petersen (4) | EMtrace | Laptop (8) | 4.2 min (1.4–8.8) | 0.19 | 7.0 |
| | EMtrace (chain2) | Laptop (8) | 5.1 min (1.3–11.6) | 0.30 | 6.9 |
| | EPI2ME wf-metagenomics | HPC (8) | 0.9 min (0.8–1.0) ‡ | 0.02 | 7.7 |
| | ViMOP | Laptop, Docker | 51 min (6–90) ‡ | 3.5 | 15.5 |
| | CZ ID | cloud | n.m. | n.m. | n.m. |
| | MEGAN-LR | — | n.r. | n.r. | n.r. |
| | INSaFLU-TELEVIR | public web server | n.m. | n.m. | n.m. |
| SARS-CoV-2 clinical, Gauthier (16) | EMtrace | Laptop (8) | 5.6 min (2.1–39.5) | 0.27 | 8.1 |
| | EPI2ME wf-metagenomics | HPC (8) | 0.8 min (0.7–1.4) ‡ | 0.01 | 7.7 |
| | ViMOP | Puhti (24) | 3.3 h (1.5–4.1) ‡ | 1.28 | 16.7 |
| | CZ ID | cloud | n.m. | n.m. | n.m. |
| | MEGAN-LR | — | n.r. | n.r. | n.r. |
| | INSaFLU-TELEVIR | public web server | n.m. | n.m. | n.m. |
| Viral mock, Buddle DNA (4) | EMtrace | HPC (24) | 39.2 min (37.5–47.1) | 6.98 | 19.1 |
| | EPI2ME wf-metagenomics | HPC (8) | 4.5 min ‡ | 0.17 | 7.8 |
| Viral mock, Buddle RNA (4) | EMtrace | HPC (24) | 29.1 min (27.6–32.1) | 4.73 | 18.0 |
| | EPI2ME wf-metagenomics | HPC (8) | 3.6 min (3.2–5.0) ‡ | 0.16 | 7.8 |
| **16S rRNA** | | | | | |
| ZymoBIOMICS mock (1; 100,128 reads) | EMtrace (Emu reference) | Laptop (8) | 45.3 min | 5.53 | 3.4 |
| | Emu | Laptop (8) | 45.0 min | 5.83 | 2.6 |
| | Spaghetti (SILVA 138) § | Laptop (8) | 18.3 min | 2.29 | 1.8 |
| | Spaghetti (Emu reference) § | Laptop (8) | 46.8 min | 6.07 | 1.7 |

| Dataset (samples) | Tool | Hardware (threads) | Wall time per sample, median (range) | CPU-hours per sample, median | Peak RAM, max (GB) |
|---|---|---|---|---|---|
| | Kraken 2 + Bracken | Laptop (8) | < 0.1 min | < 0.01 | 0.6 |
| Synthetic gut community (1; 521,661 reads) | EMtrace (Emu reference) † | Laptop (8) | 69.7 min | 7.79 | 9.6 |
| | Emu | Laptop (8) | 79.8 min | 10.1 | 5.5 |
| | Spaghetti (SILVA 138) § | Laptop (8) | 27.0 min | 3.39 | 3.4 |
| | Spaghetti (Emu reference) § | Laptop (8) | 67.1 min | 8.64 | 3.4 |
| | Kraken 2 + Bracken | Laptop (8) | 0.1 min | 0.02 | 0.7 |
| Gut Microbiome Standard, R10.4.1 (4; 72,830–84,233 reads) | EMtrace (Emu reference) | Laptop (8) | 19.8 min (18.4–21.4) | 2.37 | 3.4 |
| | Emu | Laptop (8) | 22.5 min (19.6–23.4) | 2.89 | 2.4 |
| | Spaghetti (SILVA 138) § | Laptop (8) | 15.7 min (14.3–16.9) | 2.00 | 1.8 |
| | Spaghetti (Emu reference) § | Laptop (8) | 21.1 min (19.6–21.6) | 2.63 | 1.5 |
| | Kraken 2 + Bracken | Laptop (8) | < 0.1 min | < 0.01 | 0.6 |
| | wf-16s | Laptop (8), Docker | 8.2 min (7.9–23.1) | n.m. ¶ | n.m. ¶ |

**Supplementary Table 7 | Runtime and memory per tool and dataset, as measured.** The tools ran on different hardware, so values compare tools only within a hardware class. Laptop: Apple M2 Pro, 12 cores, 16 GB RAM. HPC: CSC Roihu CPU nodes (2 × 192-core AMD EPYC 9965, Zen 5, 768 GiB RAM per node). Puhti: CSC's earlier Linux/Unix CPU supercluster, 24 cores per ViMOP run, each running at 2.1 GHz. ViMOP ran on Puhti, and on the field libraries on the laptop in Docker with amd64 emulation. The number of threads given to each run is in brackets. Ranges are over the samples of a dataset; 16S rows with one library give that run. ‡ Nextflow pipelines run with the local executor on a shared node: the value is the summed run time of the pipeline's tasks (Nextflow `realtime`), not wall time, because the recorded wall time includes waiting for CPU slots taken by concurrent runs. Peak RAM is the largest single task. † Chopper's quality filter and quality trimming off: the copy of SRR14326315 used on the laptop carries synthetic quality scores (SRA Lite); the 1,200–1,800 bp length filter was kept. § Spaghetti with its chimera-removal step skipped (Methods). ¶ wf-16s ran its tools in Docker containers, whose CPU time and memory were not recorded; the first replicate includes container start-up. n.m., not measurable (cloud service or public web server); n.r., not recorded.

### Supplementary Figure 1 | Genome recovery from real libraries

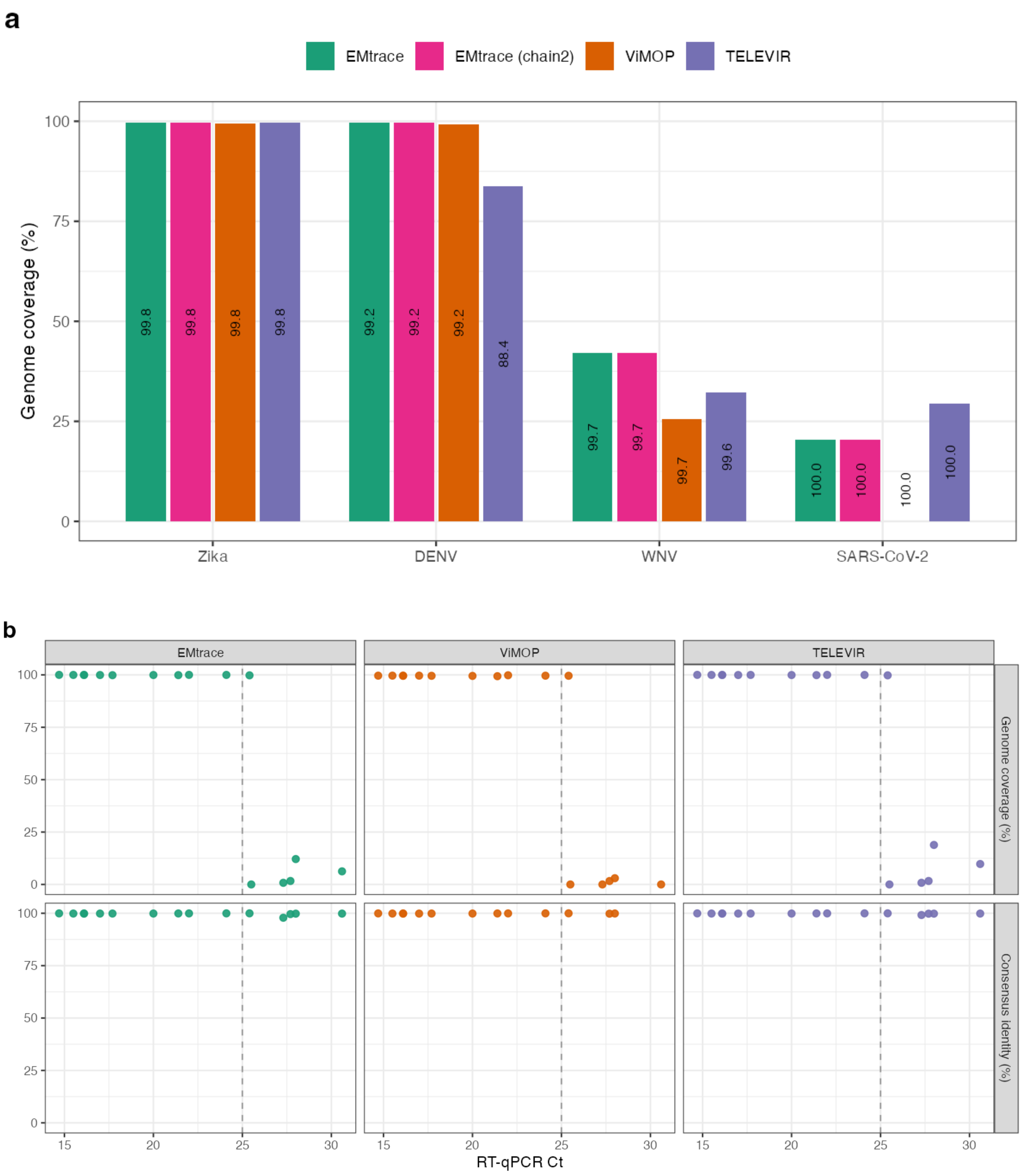


**Supplementary Figure 1 | Genome recovery from real libraries.** (a) Coverage (bar height) and identity (printed in bar) for the four field libraries; EMtrace consensus polished with Medaka. (b) Coverage (top) and identity (bottom) against RT-qPCR Ct for 16 SARS-CoV-2 samples. The dashed line marks the recovery cliff at Ct ≈ 25.

### Supplementary Note 1 | Host-filter alignment-score threshold on simulated reads

EMtrace removes host and background reads by minimap2 alignment to one or more genomes, discarding a read when its alignment score reaches a threshold (Methods). The threshold sets a trade-off: too low and genuine viral reads are discarded with the host, too high and host reads survive into the annotation step, where they cost time and can produce spurious hits. We chose the default by sweeping it on simulated reads, where every read's true origin is known.

**Data.** Simulated Nanopore reads produced with PBSIM3, the generator used for the divergence benchmark (Methods), combining the four viral read sets (SARS-CoV-2, dengue virus 2, and Lassa virus L and S segments) with simulated human and bacterial background. Each read carries a truth label giving its true origin — virus, bacterium or eukaryote — so removal can be scored per category rather than in aggregate. The sweep used viral read sets at 0, 10 and 20% divergence from their source genomes, to test whether divergent viral reads behave differently.

**Sweep.** Alignment-score thresholds of 0, 30, 50, 75, 100 and 125, with minimap2's `map-ont` seeding and, additionally, the sensitive `-k 11 -w 5` seeding. The metric is the fraction of reads removed in each truth category.

**Result.** At a threshold of **100**, 95.7% of human reads were removed, together with 0.002% of bacterial reads and no viral reads. Host removal was almost independent of the exact threshold at 100 and below, so little is gained by lowering it, while at thresholds of 50 and below a meaningful fraction of viral reads was removed as well. Viral divergence had only a small effect on viral read removal, and none above a threshold of 75: random substitutions are unlikely to create spurious matches to the human genome, at least for the viruses tested. A threshold of 100 is therefore the EMtrace default, and was used for every benchmark in this study.

The 4.3% of human reads that survive are carried into annotation, where they are classified rather than silently retained; filtering is not relied on to remove host sequence completely.
